# Forecasting range shifts of a dioecious plant species under climate change

**DOI:** 10.1101/2024.09.25.614924

**Authors:** Jacob K. Moutouama, Aldo Compagnoni, Tom E.X. Miller

## Abstract

Global warming has triggered an urgent need for predicting the reorganization of Earth’s biodiversity. Currently, the vast majority of models used to forecast population viability and range shifts in response to climate change ignore the complication of sex structure, and thus the potential for females and males to differ in their sensitivity to climate drivers. We developed demographic models of range limitation, parameterized from geographically distributed common garden experiments, with females and males of a dioecious grass species (*Poa arachnifera*) throughout and beyond its range in the south-central U.S. Female-dominant and two-sex model versions both predict that climate change will alter population viability and will induce a poleward niche shift beyond current northern limits. However, the magnitude of niche shift was underestimated by the female-dominant model, because females have broader temperature tolerance than males and become mate-limited under female-biased sex ratios. Our result illustrate how explicit accounting for both sexes could enhance population viability forecasts and conservation planning for dioecious species in response to climate change.

## Introduction

Rising temperatures and extreme drought events associated with global climate change are leading to increased concern about how species will become redistributed across the globe under future climate conditions (Bertrand et al., 2011; Gamelon et al., 2017; Smith et al., 2024). Species’ range limits, when not driven by dispersal limitation, should generally reflect the limits of the ecological niche (Lee-Yaw et al., 2016). Niches and geographic ranges are often limited by climatic factors including temperature and precipitation (Sexton et al., 2009). Therefore, any substantial changes in the magnitude of these climatic factors could impact population viability, with implications for range expansions or contractions based on which regions of a species’ range become more or less suitable (Davis and Shaw, 2001; Pease et al., 1989).

Forecasting range shifts for dioecious species (most animals and ca. 7% of plant species) is complicated by the potential for sexual niche differentiation, i.e. distinct responses of females and males to shared climate drivers (Hultine et al., 2016; Morrison et al., 2016; Pottier et al., 2021; Tognetti, 2012). For instance, the lower cost of reproduction for one sex (male or female) may allow that sex to invest its energy toward other functions that that result in higher growth rates, greater clonality, or even improved survival rates compared to the other sex, leading to sexual niche differentiation (Bruijning et al., 2017). Accounting for sexual niche differentiation is a long-standing challenge in accurately predicting which sex will successfully track environmental change and how this will impact population viability and range shifts (Gissi et al., 2023; Jones et al., 1999). Populations in which males are rare under current climatic conditions could experience low reproductive success due to sperm or pollen limitation that may lead to population decline in response to climate change that disproportionately favors females (Eberhart-Phillips et al., 2017). In contrast, climate change could expand male habitat suitability (e.g. upslope movement), which might increase seed set for mate-limited females and favor range expansion (Petry et al., 2016). Across dioecious plants, for example, studies suggest that future climate change toward hotter and drier conditions may favor male-biased sex ratios (Field et al., 2013; Hultine et al., 2016). Although the response of species to climate warming is an urgent and active area of research, few studies have disentangled the interaction between sex and climate drivers to understand their combined effects on population dynamics and range shifts, despite calls for such an approach (Gissi et al., 2023; Hultine et al., 2016).

The vast majority of theory and models in population biology, including those used to forecast biodiversity responses to climate change, ignore the complication of sex structure (but see Ellis et al., 2017; Gissi et al., 2024; Pottier et al., 2021). Traditional approaches instead focus exclusively on females, assuming that males are in sufficient supply as to never limit female fertility. In contrast, “two-sex” models are required to fully account for demographic differences between females and males and sex-specific responses to shared climate drivers (Gerber and White, 2014; Miller et al., 2011). Sex differences in maturation, reproduction, and mortality schedules can generate skew in the operational sex ratio (OSR; sex ratio of individuals available for mating) even if the birth sex ratio is 1:1 (Eberhart-Phillips et al., 2017; Shelton, 2010). Climate and other environmental drivers can therefore influence the OSR via their influence on sex-specific demographic rates. In a two-sex framework, demographic rates both influence and respond to the OSR in a feedback loop that makes two-sex models inherently nonlinear and more data-hungry than corresponding female-dominant models. Given the additional complexity and data needs, forecasts of range dynamics for dioecious species under future climate change that explicitly account for females, males, and their inter-dependence are limited (Lynch et al., 2014; Petry et al., 2016).

Tracking the impact of climate change on population viability (*λ*) and distributional limits of dioecious taxa depends on our ability to build mechanistic models that take into account the spatial and temporal context of sex specific response to climate change, while accounting for sources of uncertainty (Davis and Shaw, 2001; Evans et al., 2016). Structured population models built from demographic data collected from geographically distributed observations or common garden experiments provide several advantages for studying the impact of climate change on species’ range shifts (Merow et al., 2017; Schultz et al., 2022; Schwinning et al., 2022). First, demographic models link individual-level life history events (mortality, development, and regeneration) to population demography, allowing the investigation of factors explaining vital rate responses to environmental drivers (Dahlgren et al., 2016; Ehrlén and Morris, 2015; Louthan et al., 2022). Second, demographic models have a natural interface with statistical estimation of individual-level vital rates that provide quantitative measures of uncertainty and isolate different sources of variation, features that can be propagated to population-level predictions (Elderd and Miller, 2016; Ellner et al., 2022). Finally, structured demographic models can be used to identify which aspects of climate are the most important drivers of population dynamics. For example, Life Table Response Experiments (LTRE) built from structured models have become widely used to understand the relative importance of covariates in explaining variation in population growth rate (Czachura and Miller, 2020; Ellner et al., 2016; Hernández et al., 2023).

In this study, we combined geographically-distributed common garden experiments, hierarchical Bayesian statistical modeling, two-sex population projection modeling, and climate back-casting and forecasting to understand demographic responses to climate change and their implications for past, present, and future range dynamics. Our work focused on the dioecious plant Texas bluegrass (*Poa arachnifera*), which is distributed along environmental gradients in the south-central U.S. corresponding to variation in temperature across latitude and precipitation across longitude (Fig. S-1A). This region has experienced rapid climate warming since 1900 and this is projected to continue through the end of the century (Fig. 1). Our previous study showed that, despite evidence for differentiation of climatic niche between sexes, the female niche mattered the most in driving longitudinal range limits of Texas bluegrass (Miller and Compagnoni, 2022b). However, that study used a single proxy variable (longitude) to represent environmental variation related to aridity and did not consider variation in temperature, which is the much stronger dimension of forecasted climate change in this region (Fig. S-3). Developing a rigorous forecast for the implications of future climate change requires that we transition from implicit to explicit treatment of multiple climate drivers, as we do here. Leveraging the power of Bayesian inference, we take a probabilistic view of past, present, and future range limits by quantifying the probability of population viability (*Pr*(*λ*≥ 1)) in relation to climate drivers of demography, an approach that fully accounts for uncertainty arising from multiple sources of estimation and process error. Specifically, we asked:

1. What are the sex-specific vital rate responses to variation in temperature and precipitation across the species’ range?
2. How do sex-specific vital rates combine to determine the influence of climate variation on population growth rate (*λ*)?
3. What is the impact of climate change on operational sex ratio throughout the range?
4. What are the likely historical and projected dynamics of the Texas bluegrass geographic niche and how does accounting for sex structure modify these predictions?

**Figure 1:**
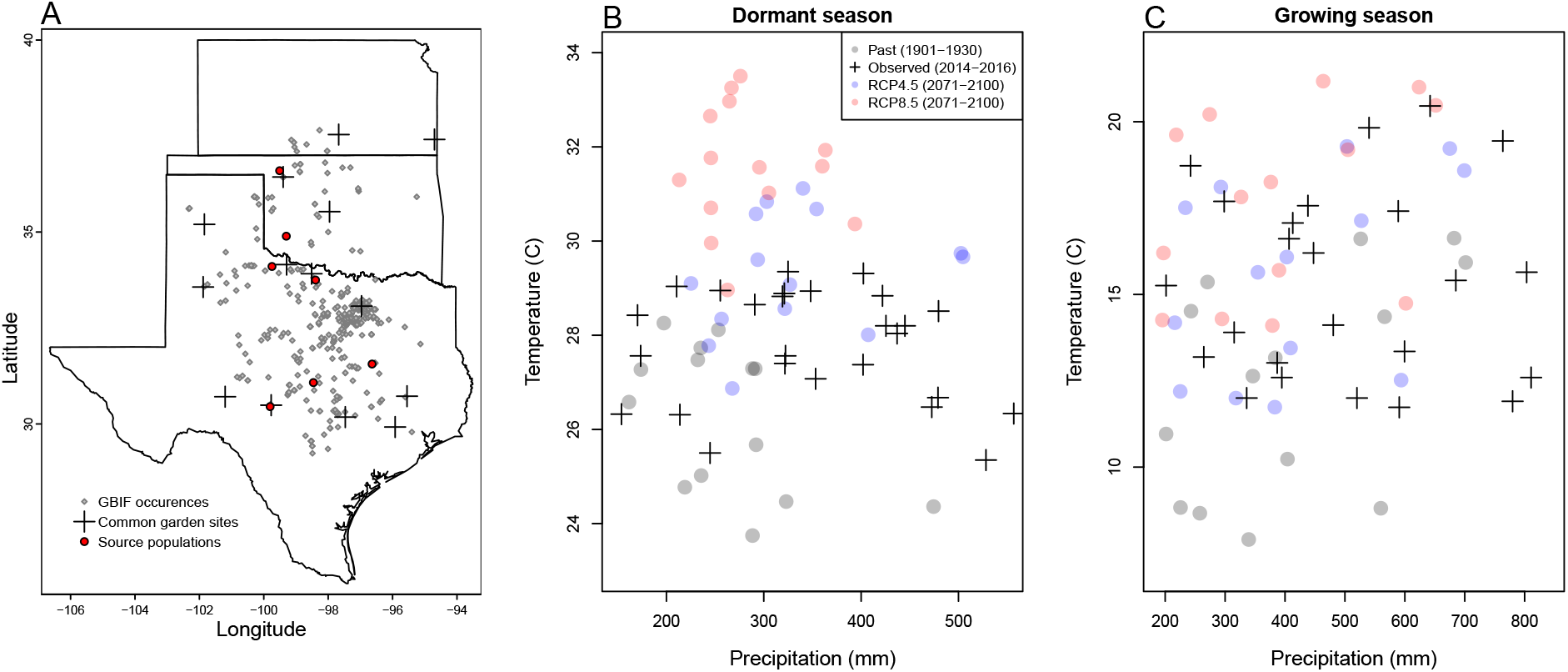
Experimental gardens and climate of the study region. **A**: Map of 14 experimental garden sites (crosses) in Texas, Oklahoma, and Kansas relative to GBIF occurrences of *Poa arachnifera* (gray points). Red points indicate source populations for plants used in the common garden experiment. **B**,**C**: Past, future, and observed climate space for growing and dormant seasons. Crosses show observed conditions for the sites and years of the common garden experiment. Gray points show historical (1901-1930) climate normals, and blue and red points show end-of-century (2071-2100) climate normals for RCP4.5 and RCP8.5 projections, respectively, form MIROC5.

## Materials and methods

### Study species and climate context

Texas bluegrass (*Poa arachnifera*) is a dioecious perennial, summer-dormant cool-season (C3) grass that occurs in the south-central U.S. (Texas, Oklahoma, and southern Kansas) (Figure 1) (Hitchcock, 1971). Texas bluegrass grows between October and May, flowers in spring, and goes dormant during the hot summer months of June to September (Kindiger, 2004). Following this life history, we divided the calendar year into growing (October 1 - May 31) and dormant (June 1 - September 30) seasons in the analyses below. Biological sex is genetically based and the birth (seed) sex ratio is 1:1 (Renganayaki et al., 2005). Females and males are morphologically indistinguishable except for their inflorescences. Like all grasses, this species is wind pollinated (Hitchcock, 1971) and most male-female pollen transfer occurs within 10-15m (Compagnoni et al., 2017). Surveys of 22 natural populations throughout the species’ distribution indicated that operational sex ratio (the female fraction of inflorescences) ranged from 0.007 to 0.986 with a mean of 0.404 (Miller and Compagnoni, 2022b).

Latitudinal limits of the Texas bluegrass distribution span 7.74 °C to 16.94 °C of temperature during the dormant season and 24.38 °C to 28.80 °C during the dormant season. Longitudinal limits span 244.9 mm to 901.5 mm of precipitation during the growing season and 156.3 mm to 373.3 mm. This region has experienced *ca*. 0.5 °C of climate warming since 1900, with faster warming during the cool-season months (0.0055°*C*/*yr*) than the hot summers (0.0046°*C*/*yr*) (Fig. S-2). Future warming is projected to accelerate to 0.03 − 0.06°*C*/*yr* by the end of the century depending on the season and forecast model. On the other hand, precipitation has increased over the past century for much of the region but is forecasted to decline back to early-20th century levels (Fig. S-2).

### Common garden experiment

#### Experimental design

We conducted a range-wide common garden experiment to quantify sex-specific demographic responses to climate variation. Details of the experimental design are provided in Miller and Compagnoni (2022b); we provide a brief overview here. The experiment was installed at 14 sites throughout and, in some cases, beyond the natural range of Texas bluegrass that sampled a broad range of latitude and longitude (Figure 1A). At each site, we established 14 blocks. For each block we planted three female and three male individuals that were clonally propagated from females and males from eight natural source populations (Figure 1A); because sex is genetically-based, clones never deviated from their expected sex. The experiment was established in November 2013 with a total of 588 female and 588 male plants, and was censused in May of 2014, 2015, and 2016. At each census, we collected data on survival, size (number of tillers), and number of panicles (reproductive inflorescences). For the analyses that follow, we focus on the 2014-15 and 2015-16 transition years, since the start of the experiment did not include the full 2013-14 transition year.

#### Climatic data collection

We gathered downscaled monthly temperature and precipitation for each site from Chelsa (Karger et al., 2017) to describe observed climate conditions during our study period. These climate data were used as covariates in vital rate regressions. We aligned the climatic years to match demographic transition years (June 1 – May 31) and growing and dormant seasons within each year. To back-cast and forecast demographic responses to changes in climate throughout the study region, we also gathered projection data for three 30-year periods: “past” (1901-1930), “current” (1990-2019) and “future” (2070-2100). Data for future climatic periods were downloaded from four general circulation models (GCMs) selected from the Coupled Model Intercomparison Project Phase 5 (CMIP5): Model for Interdisciplinary Research on Climate (MIROC5), Australian Community Climate and Earth System Simulator (ACCESS1-3), Community Earth System Model (CESM1-BGC), Centro Euro-Mediterraneo sui Cambiamenti Climatici Climate Model (CMCC-CM). All the GCMs were also downloaded from Chelsa (Sanderson et al., 2015). We evaluated future climate projections from two scenarios of representative concentration pathways (RCPs): RCP4.5, an intermediate-to-pessimistic scenario assuming a radiative forcing amounting to 4.5 *Wm*^−2^ by 2100, and RCP8.5, a pessimistic emission scenario which projects a radiative forcing of 8.5 *Wm*^−2^ by 2100 (Schwalm et al., 2020; Thomson et al., 2011).

Projection data for the three 30-year periods included warmer or colder conditions than observed in our experiment, so extending our inferences to these conditions required some extrapolation. However, across all sites, both study years were 1-2°C warmer than their corresponding “current” (1990-2019) temperature normals (Fig. S-3). Additionally, the 2014–15 growing season was generally wetter and cooler across the study region than 2015–16 (Fig. S-3). Combined, the geographic and inter-annual replication of the common garden experiment provided good coverage of most past and future conditions throughout the study region (Fig. 1B,C).

#### Sex-specific demographic responses to climatic variation across common garden sites

We used individual-level measurements of survival, growth (change in number of tillers), flowering, and number of panicles (conditional on flowering) to develop Bayesian mixed effect models describing how each vital rate varies as a function of sex, size, and four climate covariates (precipitation and temperature of growing and dormant season). These vital rate models included main effects of size (the natural log of tiller number), sex, and seasonal climate covariates (Supplementary Method S.2.1).

#### Sex ratio responses to climatic variation across common garden sites

We also used the experimental data to investigate how climatic variation across the range influenced sex ratio and operational sex ratio of the common garden populations. To do so, we developed two Bayesian linear models using data collected during three years. Each model had OSR or SR as response variable and a climate variable (temperature and precipitation of the growing season and dormant season) as predictor (Supplementary Method S.2.2).

#### Model-fitting procedures

All models were fit using Stan (Stan Development Team, 2023) in R 4.3.1 (R Core Team, 2023). We centered and standardized all climatic predictors to mean zero, variance one, which facilitated model convergence. We ran three chains for 1000 samples for warmup and 4000 for sampling, with a thinning rate of 3. We assessed the quality of the models using posterior predictive checks (Piironen and Vehtari, 2017) (Fig. S-4).

### Two-sex and female-dominant matrix projection models

We used the climate-dependent vital rate regressions estimated above, combined with additional data sources, to build female-dominant and two-sex versions of a climate-explicit matrix projection model (MPMs) structured by the discrete state variables size (number of tillers) and sex. The female-dominant and two-sex versions of the model both allow for sex-specific response to climate and differ only in the feedback between operational sex ratio and seed fertilization. For clarity of presentation we do not explicitly include climate-dependence in the notation below, but the following model was evaluated over variation in seasonal temperature and precipitation.

Let *F*_*x, t*_ and *M*_*x, t*_ be the number of female and male plants of size *x* in year *t*, where *x*∈ [1,… *U*]. The minimum possible size is one tiller and *U* is the 95th percentile of observed maximum size (35 tillers). Let 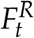 and 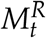 be new female and male recruits in year *t*, which we treat as distinct from the rest of the size distribution because we assume they do not reproduce in their first year, consistent with our observations. For a pre-breeding census, the expected numbers of recruits in year *t*+1 is given by:

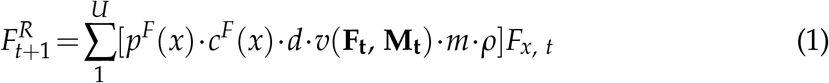

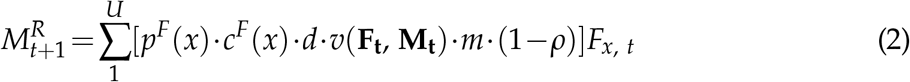

where *p*^*F*^ and *c*^*F*^ are flowering probability and panicle production for females of size *x, d* is the number of seeds per female panicle, *v* is the probability that a seed is fertilized, *m* is the probability that a fertilized seed germinates, and *ρ* is the primary sex ratio (proportion of recruits that are female), which we assume to be 0.5 (Miller and Compagnoni, 2022b).

In the two-sex model, seed fertilization is a function of population structure, allowing for feedback between vital rates and operational sex ratio (OSR). In the context of the model, OSR is defined as the fraction of panicles that are female and is derived from the *U ×* 1 vectors **F**_**t**_ and **M**_**t**_:

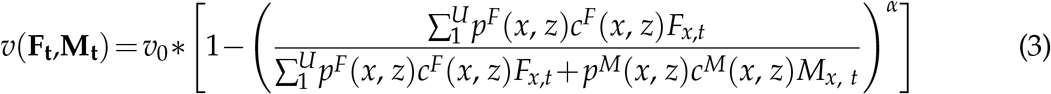

The summations tally the total number of female and male panicles over the size distribution, giving the fraction of total panicles that are female. We focus on the female fraction of panicles and not female fraction of reproductive individuals because panicle number can vary widely depending on size; we assume that few males with many panicles vs. many males with few panicles are interchangeable pollination environments. Eq. 3 has the properties that seed fertilization is maximized at *v*_0_ as OSR approaches 100% male, goes to zero as OSR approaches 100% female, and parameter *α* controls how female seed viability declines as male panicles become rare. We estimated these parameters using data from a sex ratio manipulation experiment, conducted in the center of the range, in which seed fertilization was measured in plots of varying OSR; this experiment is described elsewhere (Compagnoni et al., 2017) and is summarized in Supplementary Method S.2.3. This experiment also provided estimates for seed number per panicle (*d*) and germination rate (*m*). Lacking data on climate-dependence, we assume that seed fertilization, seed number, and germination rate do not vary with climate.

The dynamics of the size-structured component of the population are given by:

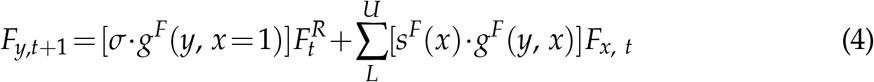

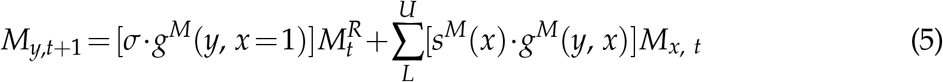

The first terms indicate recruits that survived their first year and enter the size distribution of established plants. We estimated the seedling survival probability *σ* using demographic data from the congeneric species *Poa autumnalis* in east Texas (T.E.X. Miller and J.A. Rudgers, *unpublished data*), and we assume that *σ* is the same across sexes and climatic variables. We did this because we had little information on the early life cycle transitions of greenhouse-raised transplants. We used *g* (*y, x* = 1) (the future size distribution of one-tiller plants from the transplant experiment) to give the probability that a surviving recruit reaches size *y*. The second component of the equations indicates survival and size transition of established plants from the previous year, where *s* and *g* give the probabilities of surviving at size *x* and growing from sizes *x* to *y*, respectively, and superscripts indicate that these functions may be unique to females (*F*) and males (*M*).

The model described above yields a 2(*U* + 1)*×*2(*U* + 1) transtion matrix. We estimated the population growth rate *λ* of the female dominant model as the leading eigenvalue of the transition matrix. Since the two-sex MPM is nonlinear (matrix elements affect and are affected by population structure) we estimated *λ* and stable sex ratio (female fraction of all individuals) and operational sex ratio (female fraction of panicles) by numerical simulation. Since all parameters were estimated using MCMC sampling, we were able to propagate the uncertainty in our estimates of the vital rate parameters to uncertainty in *λ*. Furthermore, by sampling over distributions associated with site, block, and source population variance terms, we are able to incorporate process error into the total uncertainty in *λ*, in addition to the uncertainty that arises from imperfect knowledge of the parameter values. For example, sampling over site and block variances accounts for regional and local spatial heterogeneity that is not explained by climate, and sampling over source population variance accounts for genetically-based demographic differences across the species’ range.

### Life Table Response Experiments

We conducted two types of Life Table Response Experiments (LTRE) to isolate contributions of climate variables and sex-specific vital rates to variation in *λ*. First, to identify which aspect of climate is most important for population viability, we used an LTRE based on a nonparametric model for the dependence of *λ* on parameters associated with seasonal temperature and precipitation (Ellner et al., 2016). To do so, we used the RandomForest package to fit a regression model with four climatic variables (temperature of growing season, precipitation of growing season, temperature of the dormant season and precipitation of the dormant season) as predictors and *λ* calculated from the two sex model as response (Liaw et al., 2002). The regression model allowed the estimation of the relative importance of each predictor.

Second, to understand how climate drivers influence *λ* via sex-specific demography, we decomposed the effect of each climate variable on population growth rate (*λ*) into contribution arising from the effect on each female and male vital rate using a “regression design” LTRE (Caswell, 1989, 2000). This LTRE decomposes the sensitivity of *λ* to climate according to:

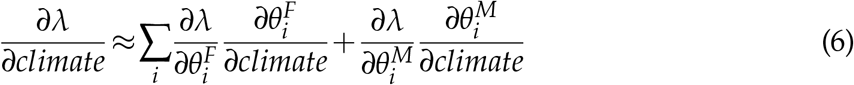

where, 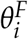 and 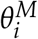 represent sex-specific parameters (the regression coefficients of the vital rate functions). Because LTRE contributions are additive, we summed across vital rates to compare the total contributions of female and male parameters.

### Population viability across the climatic niche and geographic range

To understand how climate shapes the niche and geographic range of Texas bluegrass, we estimated the probability of self-sustaining populations (Pr (*λ*≥ 1)) conditional to temperature and precipitation of the dormant and growing seasons. Pr (*λ* > 1) was calculated for the two-sex model and the female dominant MPMs using the proportion of the 300 posterior samples that lead to a *λ*≥ 1 (Diez et al., 2014). Population viability in climate niche space was then represented as a contour plot with values of Pr (*λ*> 1) at given temperature and precipitation for the growing season, holding dormant season climate constant, and vice versa.

Pr (*λ*> 1) was also mapped onto geographic layers of three US states (Texas, Oklahoma and Kansas) to delineate past, current and future potential geographic distribution of the species. To do so, we estimated Pr (*λ* > 1) conditional to all climate covariates for each pixel (∼25 km2) for each time period (past, present, future). Because of the amount of the computation involved, we use 100 posterior samples to estimate Pr (*λ*> 1) across the study area (Texas, Oklahoma and Kansas).

## Results

### Sex specific demographic responses and sex ratio variation across climatic conditions

We found strong demographic responses to climate drivers across our Texas bluegrass common garden sites and years, and evidence for demographic differences between the sexes. Regression coefficients related to sex and/or sex:size interactions were significantly non-zero (95% credible intervals excluding zero) for most vital rates (Fig. S-5), suggesting sexual divergence in demography. Females generally had an advantage over males, especially in survival and flowering (Fig. 2). Furthermore, there were significant interactions between sex and one or more climate variables, particularly for growth (Fig. S-5B), indicating sexual niche divergence in response to shared climate drivers. Fig. S-6 and S-7 visualize the magnitude of sexual divergence in demography across niche space, revealing that female advantages in flowering and panicle production were greatest at both high and low growing season temperature extremes.

**Figure 2:**
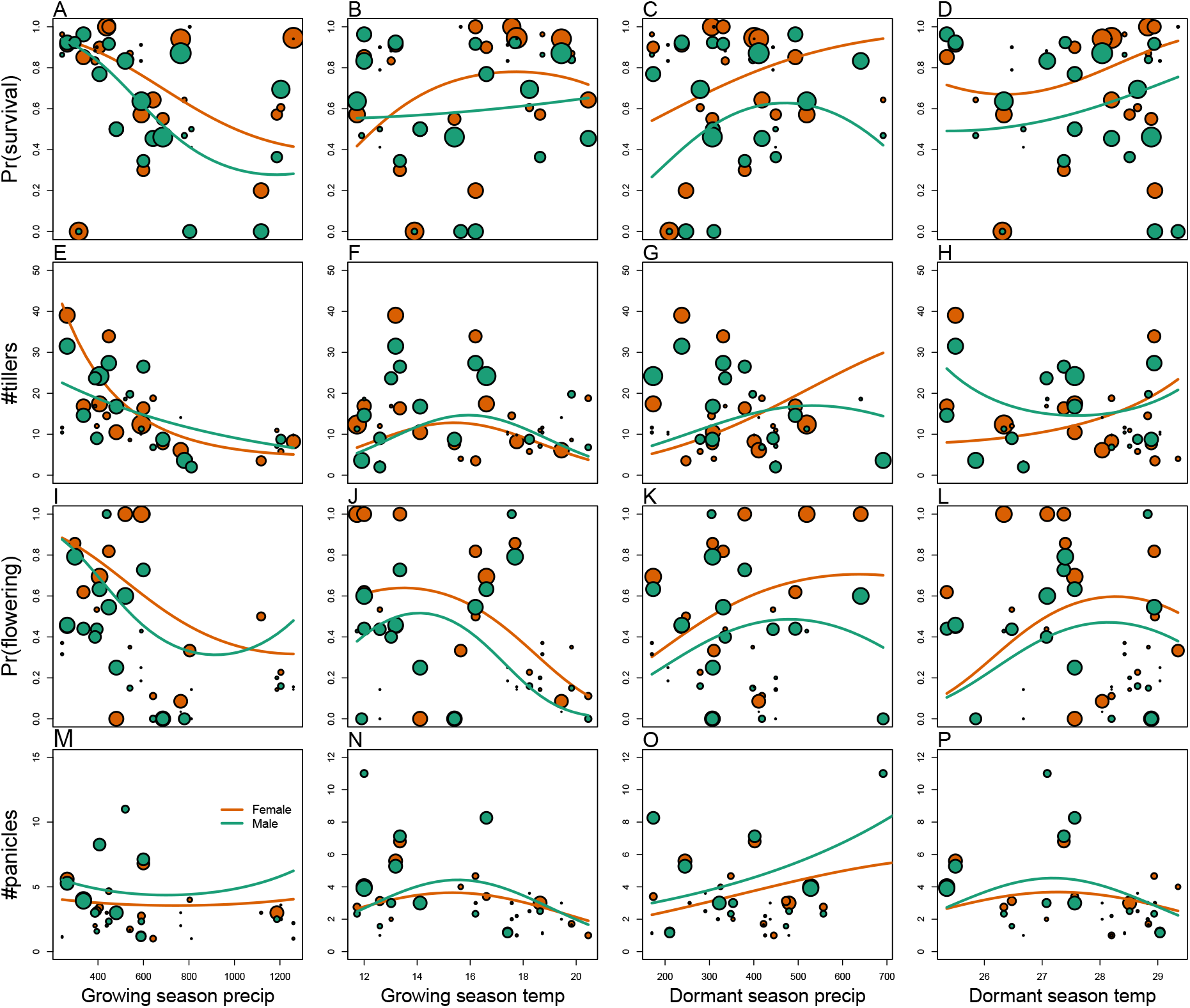
Sex specific demographic response to climate across species range. (A, B, C, D) Probability of survival as a function of precipitation and temperature of the growing and dormant season. (E, F, G, H) Change in number of tillers as a function of precipitation and temperature of the growing and dormant season. (I, J, K, L) Probability of flowering as a function of precipitation and temperature of growing and dormant season. (M, N, O, P) Change in number of panicles as a function of precipitation and temperature of the growing and dormant season. Points show means by site for females (orange) and males (green). Points sizes are proportional to the sample sizes of the mean and are jittered. Lines show fitted statistical models for females (orange) and males (green) based on posterior mean parameter values.

Across common garden sites, operational sex ratio (proportion of panicles that are female) of the experimental populations was female-biased on average (≈ 60 % female), reflecting the overall greater rates of female vs. male flowering rather than bias in the underlying population composition. OSR was most female-biased (up to 80% female) at extreme values of temperature, especially growing season temperature (Fig. S-8, Fig. S-9), consistent with the female reproductive advantage at temperature extremes seen in the vital rate data (Fig. S-6). In contrast, there was very little variation in sex ratio (proportion of plants that are female) in the years following common garden establishment (all sites were planted with equal numbers of females and males) and no detectable influence of climate covariates (Fig. S-10), indicating that skew in the OSR comes from sex-biased reproductive rates more so than sex-biased survival.

### Climate drivers of population viability across niche space

Putting all vital rates together in the MPM framework reveals how climate shapes fitness variation across niche dimensions and geographic space, and how accounting for sex structure modifies these inferences. For both female-dominant and two-sex models, fitness variation across niche space was dominated by temperature, with weaker effects of precipitation (compare vertical and horizontal contours in Fig. 3). These visual trends are supported by LTRE decomposition indicating that variation in fitness across climatic conditions is most strongly driven by responses to growing and dormant season temperature, with weaker interactive effects of precipitation that modulate the effects of temperature (Fig. S-12). LTRE analysis also showed that declines in population viability at high and low temperatures were driven most strongly by reductions in vegetative growth and panicle production, with stronger contributions from females than males (Fig. S-13). Intermediate temperatures of both growing and dormant seasons were associated with near-certain projections of population viability (*Pr*(*λ*≥ 1) ≈ 1), and high and low temperature extremes during both seasons were associated with low niche suitability (*Pr*(*λ*≥ 1) < 0.2). Higher precipitation slightly expanded the range of suitable temperatures during the dormant season (Fig. 3A), and the reverse was true in the growing season (Fig. 3B). Points in Fig. 3 show that climate change forecasted for the common garden locations would move many of them toward lower-suitabilty regions of niche space associated with high growing and dormant season temperatures (see also Fig. S-14).

**Figure 3:**
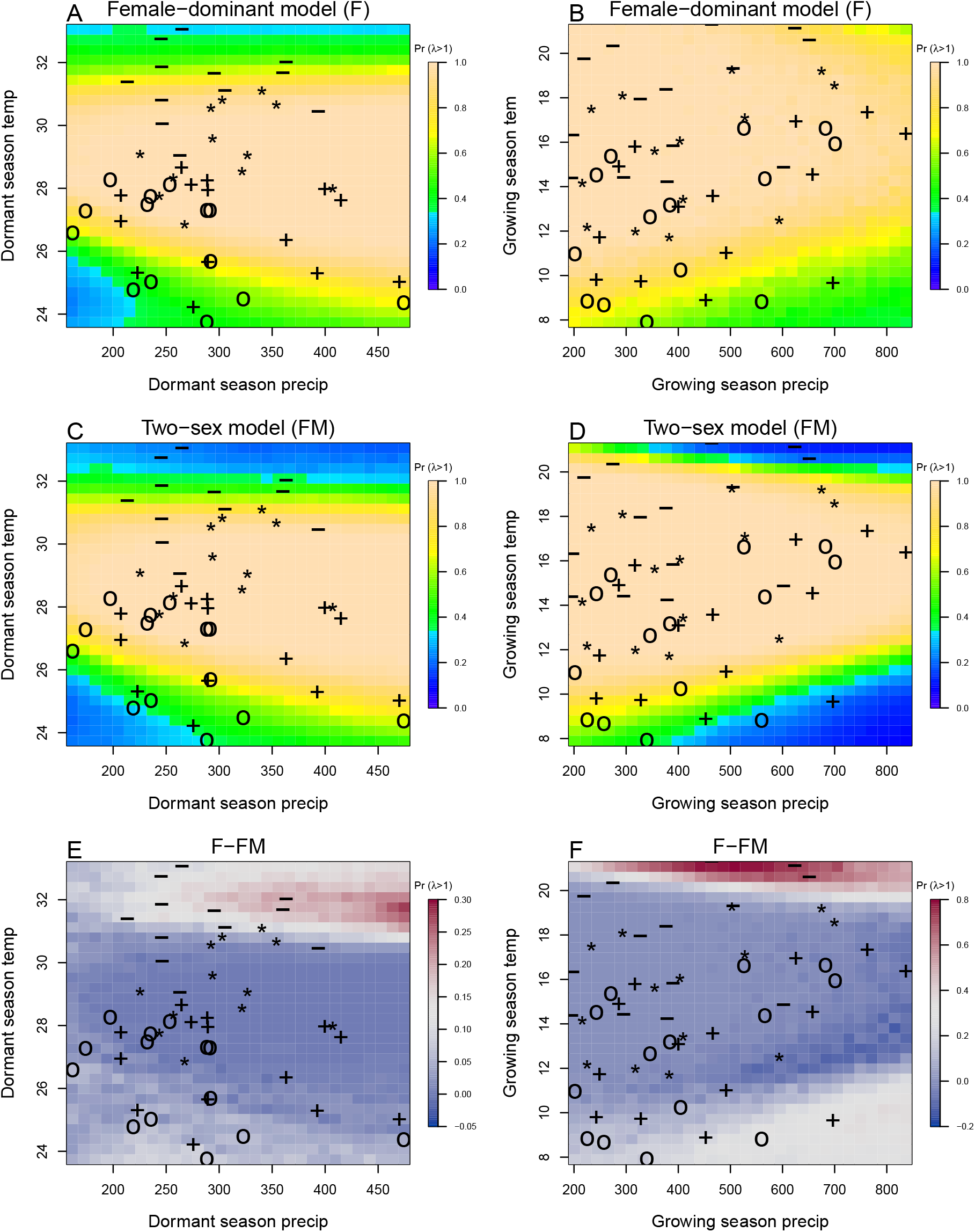
A two-dimensional representation of the predicted niche shift over time (past, present and future climate conditions). “**o**”: Past, “**+**”: Current,”**\***”: RCP 4.5,”**-**”: RCP 8.5.

While the female-dominant and two-sex models were generally in agreement about high confidence in intermediate temperature optima, they differed around the edges of niche space (Fig. 3C,D,S-14). The female-dominant model over-predicted population viability, especially with respect to growing season temperature. For example, the female-dominant model predicted that, for most levels of precipitation, warm growing season (winter) temperatures of ∼ 20°*C* had high suitability (*Pr*(*λ*≥ 1) > 0.9), while the two-sex model indicated that these conditions were most likely unsuitable (*Pr*(*λ*≥ 1) < 0.5). Similarly, at low winter temperatures that the two-sex model identifies with high certainty as unsuitable (*Pr*(*λ* ≥ 1) < 0.1), the female-dominant model is more optimistic (*Pr*(*λ*≥ 1) > 0.4). Across growing season climate space, the female-dominant model over-estimates population viability by ca. 9.23%, on average (Fig. 3D, Fig. S-15B). The difference between female-dominant and two-sex models was qualitatively similar but weaker in magnitude for niche dimensions of the dormant season (Fig. 3C, Fig. S-15A). Female-dominant and two-sex models diverged most strongly in regions of niche space that favored strongly female-biased operational sex ratios (Fig. S-16). This suggests mate limitation as the biological mechanism underlying model differences. The two-sex model accounts for feedbacks between OSR and female fertility, with reduced seed viability at OSR exceeding ∼ 75% female panicles (Fig. S-17) Lacking this feedback, the female-dominant model over-predicts population viability in regions of niche space where male flowering is not sufficient to maximize seed set.

### Climatic change induces shifts in geographic niche and population OSR

Projecting the climatic niche onto geographic space, we find that suitable niche conditions for Texas bluegrass are shifting north under climate change (Fig. 4). Under past (1901-1930) and present (1990-2019) conditions, the two-sex and female-dominant models predict widespread suitability with high confidence (*Pr*(*λ* ≥ 1) ≈ 1) across much of Texas and Oklahoma. For both models, the predicted geographic niche generally corresponds well to independent observations of the Texas bluegrass distribution (Fig. 4). The predicted geographic niche is more expansive than the GBIF occurrences, particularly at southern, western, and eastern edges, suggesting some degree of range disequilibrium (e.g., due to dispersal limitation), geographic bias in occurrence records, and/or model mis-specification. Comparing past to present conditions, the geographic niche for both models has shifted slightly poleward, with reductions in viability at the southern margins and expansions of viability at northern margins. The northward shift of suitable niche conditions is even more pronounced in projections to end-of-century (2071-2100) conditions, with the most dramatic changes in the most pessimistic (RCP8.5) scenario (Fig. 4). In fact, under the pessimistic scenario, Texas bluegrass will have very little remaining climate suitability in the state of Texas by the end of the 21st century. The predicted poleward niche shift is consistent across different global circulation models (Fig. S-22, Fig. S-23, Fig. S-24).

Female-dominant and two-sex models are in broad agreement about northward migration of the climatic niche, but the geographic projections reveal hotspots of disagreement where the female-dominant model over-predicts climate suitability and under-predicts the likelihood of range shifts (Fig. 4). These hotspots are generally regions of predicted female bias in the operational sex ratio (Fig. S-18). The strongest contrast between the two models is in the pessimistic climate change scenario (RCP8.5), where the female-dominant model over-predicts population viability by as much as 20% across much of the region (Fig. S-25) and thus under-estimates the magnitude of a potential range shift. In this scenario, a broad swath of the current distribution that is forecasted to by effectively unsuitable (*Pr*(*λ*≥ 1) ≈ 0) by the two-sex model is identified as marginally suitable (*Pr*(*λ*≥ 1) ≈ 0.5) by the female-dominant model. Accordingly, the OSR of Texas bluegrass across its range is projected to be ca. 75% female panicles, on average, by end of century under RCP8.5, an increase from ca. 60% female under projections for past and current conditions (Fig. 5). The more optimistic climate change scenario (RCP4.5) predicts an intermediate shift in OSR, with hotspots of change at northern and southern range edges becoming strongly female-biased but most of the range remaining near current levels of 60% female (Fig. 5; Fig. S-21,Fig. S-19,Fig. S-20).

**Figure 4:**
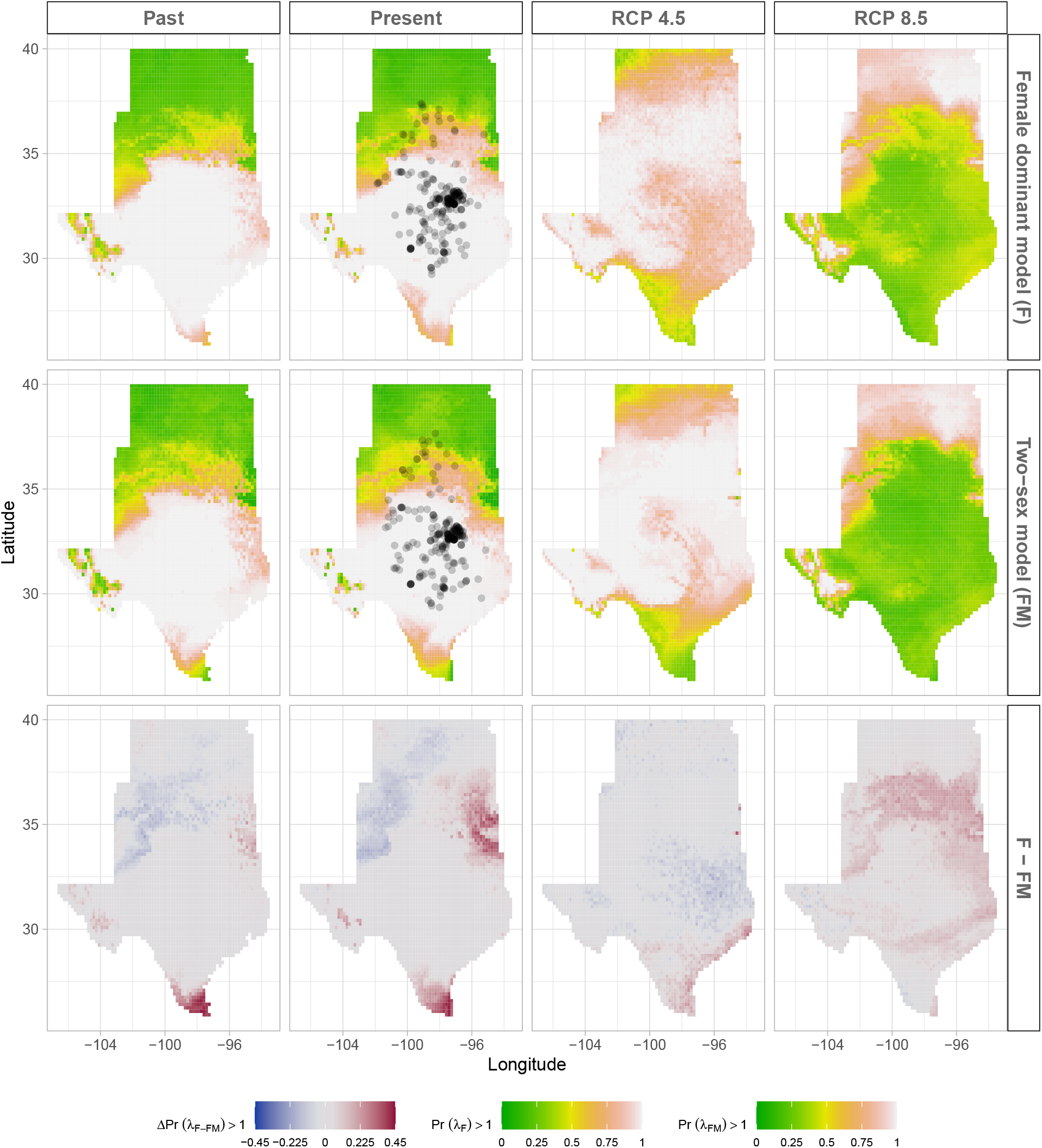
Climate change favors range shift towards north edge. (A) Past, (B) Current, (C and D) Future predicted range shift based on the predicted probabilities of self-sustaining populations, Pr (*λ*> 1), using the two-sex model that incorporates sex-specific demographic responses to climate with sex ratio dependent seed fertilization. (E) Past, (F) Current, (G and F) Future predicted range shift based on the predicted probabilities of self-sustaining populations, Pr (*λ*> 1), using the female dominant model. Future projections were based on the CMCC-CM model. The black dots on panel B and F indicate all known presence points collected from GBIF. The occurrences of GBIFs are distributed in with higher population fitness habitat Pr (*λ*> 1), confirming that our study approach can reasonably predict range shifts.

**Figure 5:**
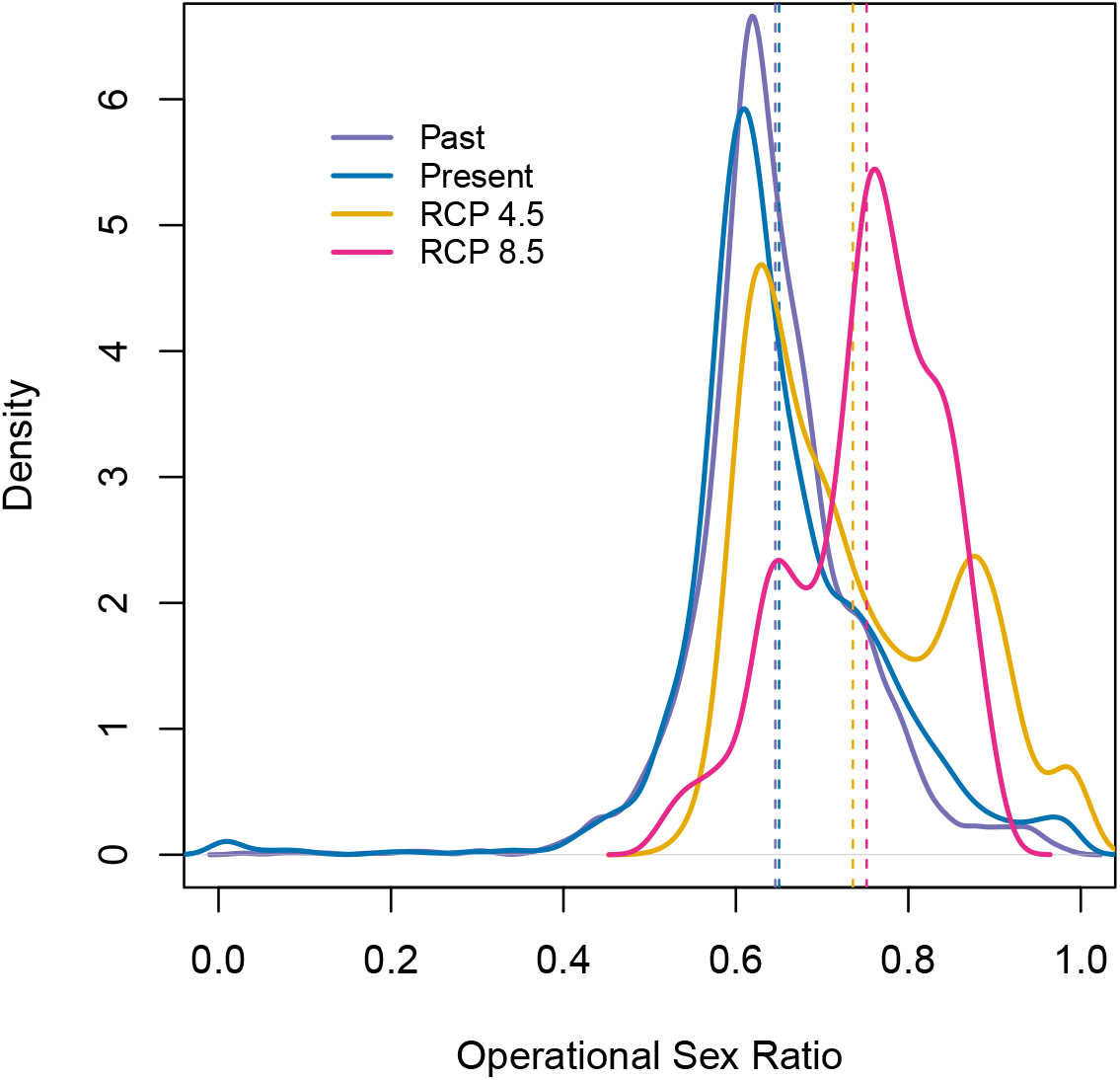
Change in Operational Sex Ratio (proportion of female panicle) over time (past, present, future). Future projections were based on the CMCC-CM model. The mean sex ratio for each time period is shown as vertical dashed line.

## Discussion

Dioecious species make up a large fraction of Earth’s biodiversity – most animals and many plants – yet we have little knowledge about how sex-specific demography and responses to climate drivers may affect population viability and range shifts of dioecious species under climate change. We used demographic data collected from common garden experiments, hierarchical Bayesian statistical modeling, and sex-structured demographic modeling to forecast for the first time the likely impact of climate change on range dynamics of a dioecious species. We found that demographic rates of Texas bluegrass and their sensitivities to climate drivers show significant sex bias, with females out-performing males, on average, and high and low temperature extremes disproportionately favoring female reproduction, leading to female skew in the operational sex ratio. In fact, we show that future climate change will likely not only shift this species’ geographic niche northward, but it will also skew operational sex ratios toward stronger female bias. Our two-sex modeling framework accounts for reductions in female fertility with increasing female bias, and therefore predicts a narrower climatic niche than the corresponding female-dominant model that ignores the feedback between population structure and vital rates. Failure to account for population sex structure can therefore lead to overestimation of suitable niche space and underestimation of range shifts under global change. Our finding that climate change in the south-central US will likely lead to female-biased operational sex ratios contrasts with previous studies of dioecious plants. While a baseline female demographic advantage has been observed in several dioecious species (Bawa, 1980; Sasaki et al., 2019; Zhao et al., 2012), focused on sex-specific sensitivity to climate drivers predict an increase in male frequency in response in climate change (Hultine et al., 2016; Petry et al., 2016). We speculate that differences in the costs of reproduction related to pollination mode may help explain which sex is favored under climate stress. For most dioecious plant species, the cost of reproduction is often higher for females than males due to the requirement to develop seeds and fruits (Hultine et al., 2016). However, several studies reported a higher cost of reproduction for males in wind pollinated species due to the larger amounts of pollen they produce (Bruijning et al., 2017; Bü rli et al., 2022; Cipollini and Whigham, 1994; Field et al., 2013). Additional comparative studies across species that differ in life history traits are needed to draw inferences regarding which types of species are likely to become female-or male-biased in response to global change stressors.

While a two-sex modeling approach clearly adds biological realism, it was also additional work (in the form of experiments, data, equations, code, and computation). Was it worth the trouble? Generally, we suggest the answer should depend on the aims of the investigator. Predictions of the sex-structured and female-dominant models were in strong agreement about climate niche optima, and LTRE decomposition suggested that female vital rates determine population responses to climate variation much more so than male vital rates. If we wanted to know whether a poleward range shift is likely for Texas bluegrass, the simpler female-dominant approach could have given us the correct answer. But more focused questions, especially around the edges of niche space where sex ratio skew is more likely to impair population viability, may require an explicit accounting for sex structure. If we aimed to identify specific regions that are more or less inclined toward contraction or expansion, or sites that might be suitable for assisted migration, we might reach qualitatively different conclusions with female-dominant and two-sex models. For example, the female-dominant model is over-confident that large swaths of Oklahoma will remain marginally suitable for Texas bluegrass under the business-as-usual emissions scenario, while the two-sex model is more pessimistic, because this region will become too female-biased to support viable populations. More generally, we hypothesize that accounting for sex structure should be most important under conditions that are already near the limits of population viability, where effects of mate limitation could be more consequential. This suggests a particularly important role of sex-structured modeling for threatened and endangered species, as conservation biologists have already recognized (Jenouvrier et al., 2012; Milner-Gulland, 1994; Molnár et al., 2010).

Our results suggest that climate change, and specifically climate warming, will drive a classic pattern of poleward expansion: contraction at the southern trailing edge due to temperatures exceeding tolerable limits and expansion at the northern leading edge due to release from low temperature limitation. Our statistical models captured temperature-dependence in a phenomenological way, and the physiological mechanisms underlying these responses remain to be explored. Increasing temperature could increase evaporative demand, affect plant phenology (Iler et al., 2019; McLean et al., 2016; Sherry et al., 2007), and germination rate (Reed et al., 2021). The potential for temperature to influence these different processes changes seasonally (Konapala et al., 2020). For example, studies suggested that species that are active during the growing season such as cool grass species can have delayed phenology in response to global warming, particularly if temperatures rise above their physiological tolerances (Cleland et al., 2007; Williams et al., 2015).

Regardless of the mechanism, it is clear that climate warming will generate leading and trailing edges. Whether and at what pace the realized species’ distribution tracks geographic changes in suitable niche space is a different, open question. Expansion of the leading edge could lag behind availability of suitable habitat due to dispersal limitation (Pagel et al., 2020), and legacies of long-lived individuals can promote persistence of trailing edge populations even as environmental conditions deteriorate (Margaret EK et al., 2023). Environmentally-explicit demographic models are emerging as powerful tools to understand and predicts the limits of population viability under global change (Merow et al., 2017; Schultz et al., 2022), but incorporating non-equilibrium dynamics that emerge from dispersal limitation and and historical legacies is an important new direction for this field.

Our forecasts for responses to climate change in Texas bluegrass should be interpreted in light of several features of our study design. First, the design of our common garden experiment and statistical modeling means that our geographic projections correspond to an “average” genotype from across the range of Texas bluegrass. Local adaptation to climate could make southern and northern edge populations more resilient to high and low temperature stress, respectively, than the range-wide average (Angert et al., 2020; Gilbert et al., 2017). The role of local adaptation in mitigating population response to climate is an important next step in forecasting species’ responses to global change. Second, as is true for many ecological systems, future climate is likely to include conditions that have no present-day analog (Intergovernmental Panel On Climate Change (Ipcc), 2023), a major challenge for ecological forecasting. The years and locations of our experiment provided us with unusually good coverage of likely past, present, and future conditions expected throughout the study region, but we still had to extrapolate the statistical models to predict responses to colder winter temperatures (that were more common in the past) and hotter summer temperatures (that are expected in the future) than we directly observed (Fig. 1). By employing a probabilistic measure of niche and geographic suitability (*Pr*(*λ*) ≥ 1), our projections account for the uncertainty associated with these extrapolated climate responses, but there would be value in combining the spatiotemporal sampling of a common garden design with experimental manipulations that push systems toward historical and/or future conditions. Third, while we incorporated uncertainty associated with parameter estimation and process error, there is additional uncertainty in future climate conditions. Future forecasts for Texas bluegrass were generally consistent across different global circulation models (reference supp figures), but combining uncertainty in future conditions alongside uncertainty in biological responses to those conditions is an important frontier in ecological forecasting (Dietze et al., 2018).

## Conclusion

We investigated how demographic differences between the sexes and contrasting sensitivity to climate can drive skewness in sex ratio on and possible range shifts in the context of climate change. For Texas bluegrass, the future is female, and it is in Kansas. Our results suggest that tracking only females could lead to an underestimation of the effect of climate change on population dynamics, because it misses the feedback between population structure and female fertility. But in broad strokes, a female-dominant perspective tells much of the story, and that will likely be true for dioecious plants and animals with mating systems in which few males can fertilize many females. Our work also provides a framework for predicting the impact of global change on population dynamics and range shifts using probabilistic measures that can incorporate and pick apart the many types of uncertainty that arise when reconstructing the past or forecasting the future.

## Acknowledgements

This research was supported by National Science Foundation Division of Environmental Biology awards 2208857 and 2225027. We thank the organizations and institutions who hosted us at their field station facilities, including The Nature Conservancy, Sam Houston State University, University of Texas, Texas A&M University, Texas Tech University, Lake Lewisville Environmental Learning Area, Wichita State University, and Pittsburgh State University.

## Supporting Information

### S.1 Supporting Figures

**Figure S-1:**
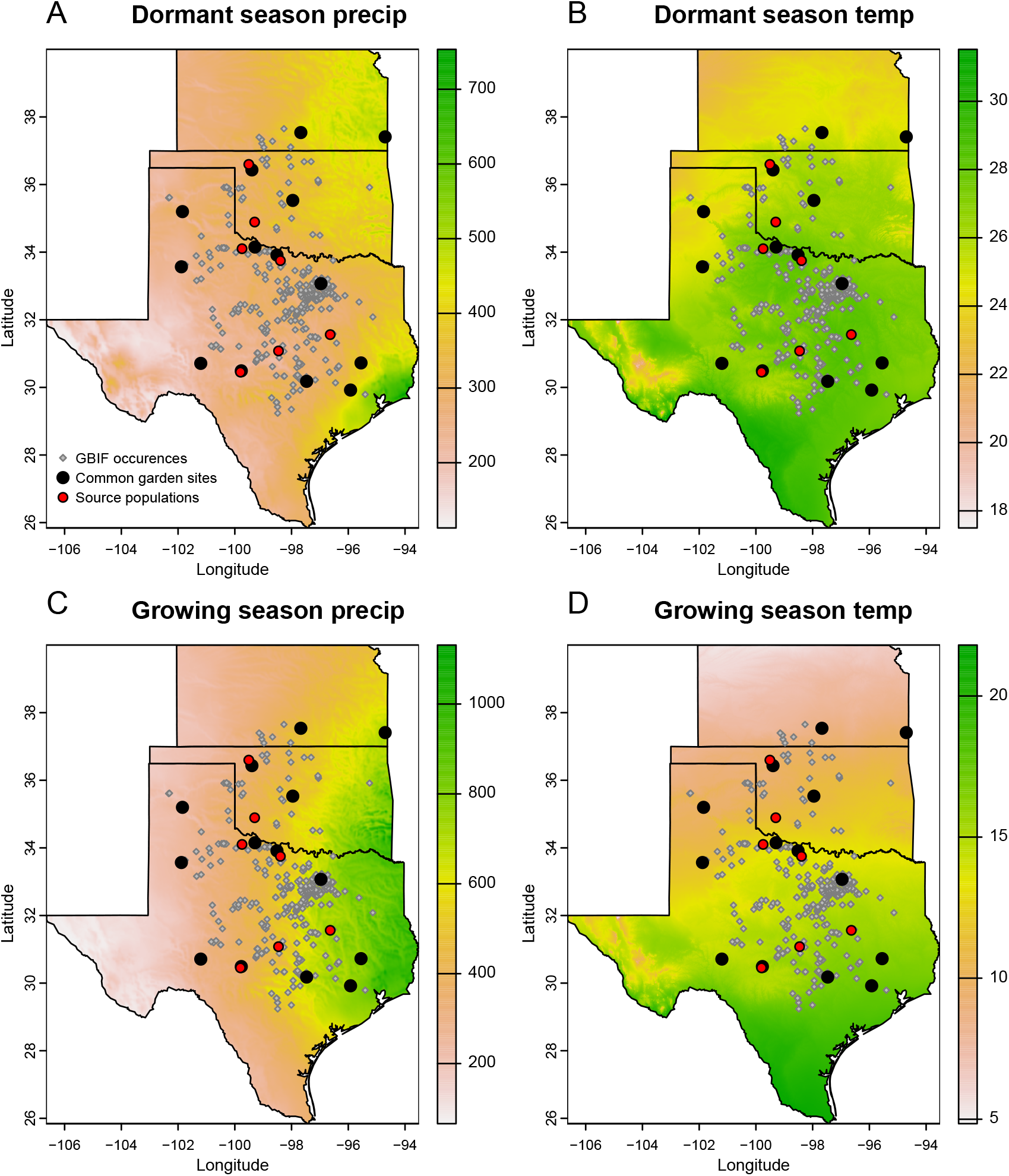
Maps of 30-year (1990-2019) climate normals and study sites used for demographic monitoring of *Poa arachnifera* in Texas, Kansas and Oklahoma in the United States. Precipitation is in mm and temperature is in °C. We surveyed 22 natural populations (grey diamond). The common garden sites were installed on 14 sites (black circle) collected from 7 sources populations (red circle).

**Figure S-2:**
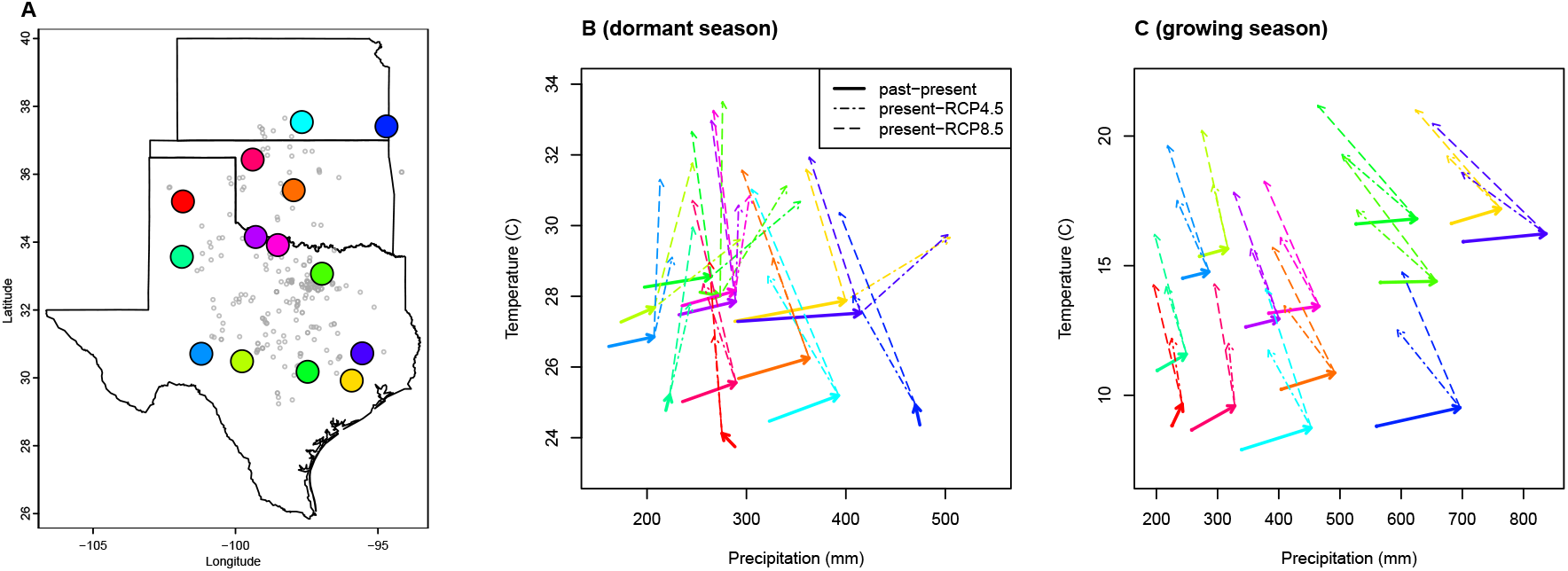
(A), Common garden locations in Texas, Oklahoma, and Kansas (points) and (B,C) corresponding changes in growing and dormant season climate. Arrows in B,C connect past (1901-1930) and present (1990-2019) climate normals, and present and future (2071-2100) climate normals under RCP4.5 and RCP8.5 forecasts. Future forecasts are from MIROC5 but other climate models show similar patterns.

**Figure S-3:**
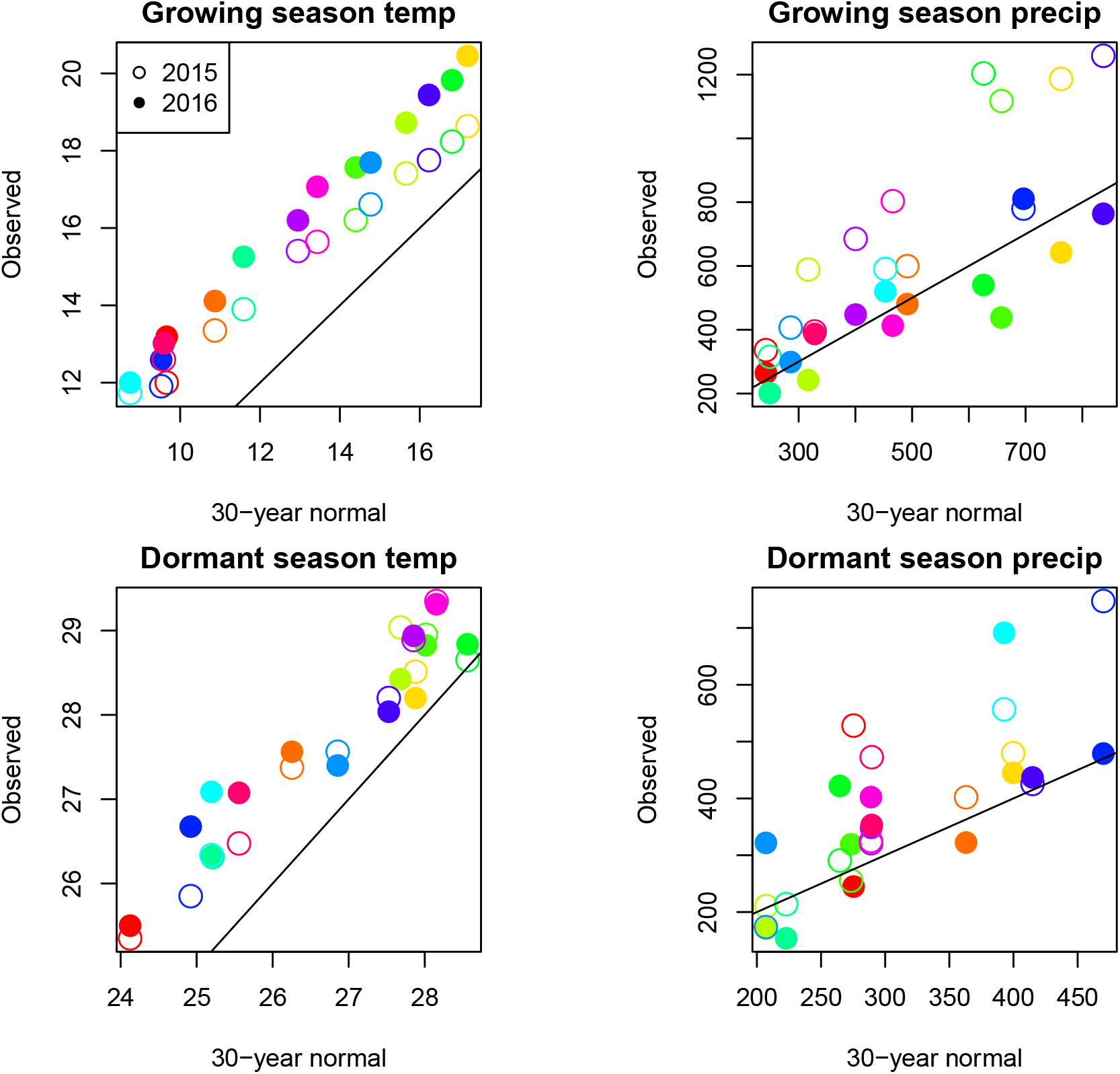
Comparison of 30-year (1990-2019) climate normals and observed weather conditions at common garden sites during the 2014–15 and 2015–16 census periods. Temperature is in °C and precipitation is in *mm*. Colors represent sites and lines show the *y* = *x* relationship.

**Figure S-4:**
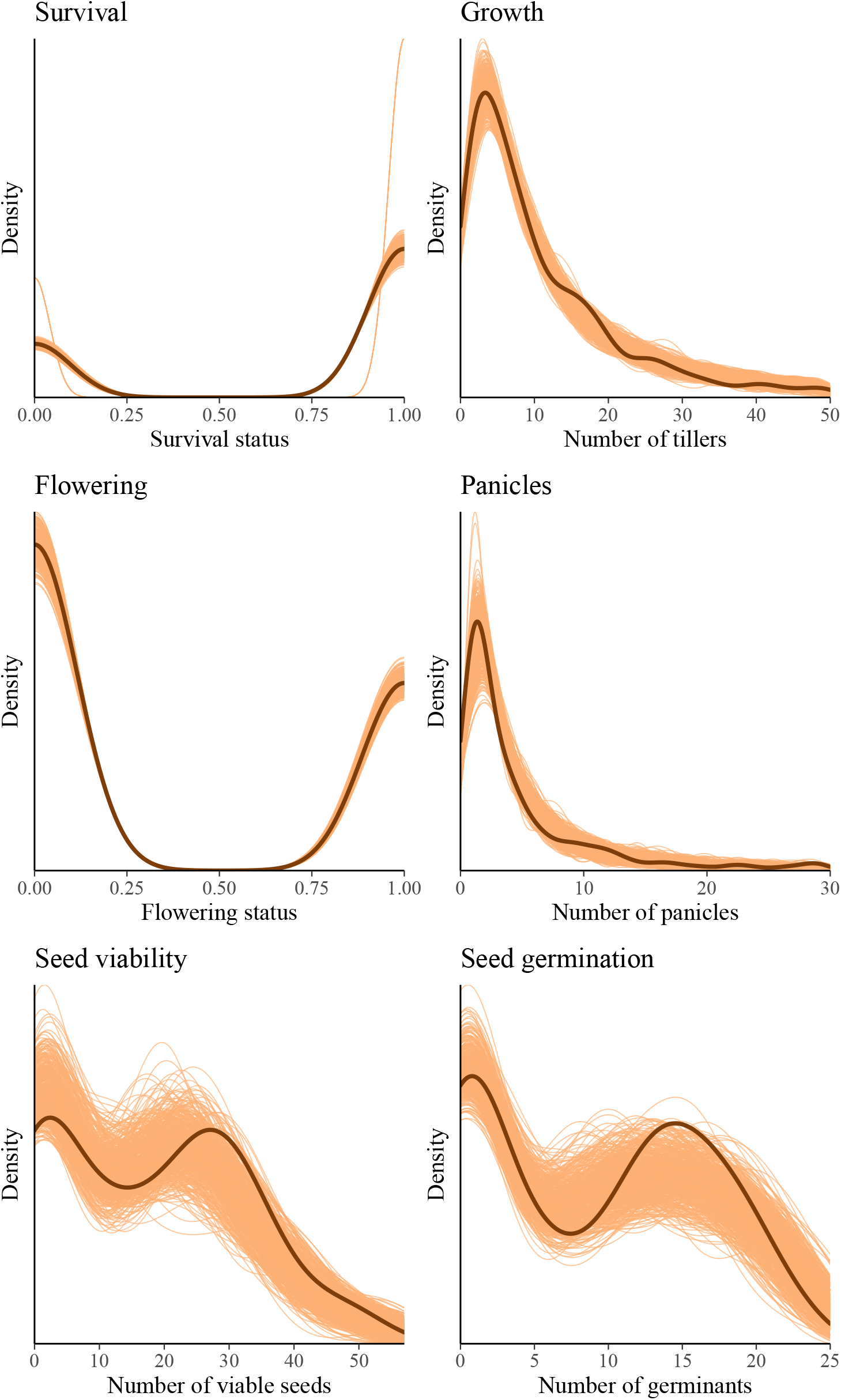
Posterior predictive checks. Consistency between real data and simulated values suggests that the fitted vital rate models accurately describes the data. Graph shows density curves for the observed data (light orange) along with the simulated values (dark orange).

**Figure S-5:**
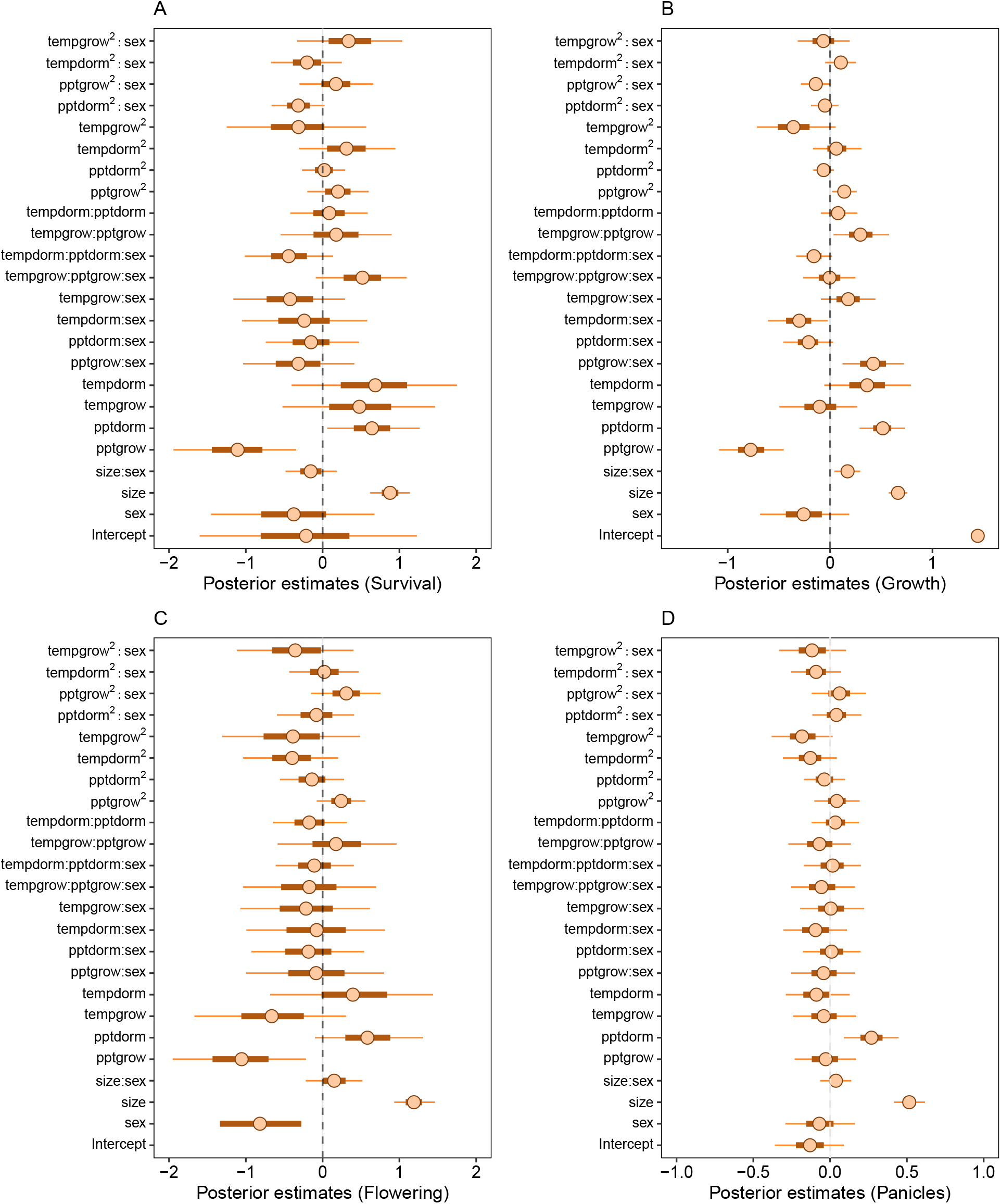
Mean parameter values and 95% credible intervals of the posterior probability distributions. pptgrow is the precipitation of growing season, Tempgrow is the temperature of growing season, pptdorm is the temperature of dormant season, Tempdorm is the temperature of dormant season.

**Figure S-6:**
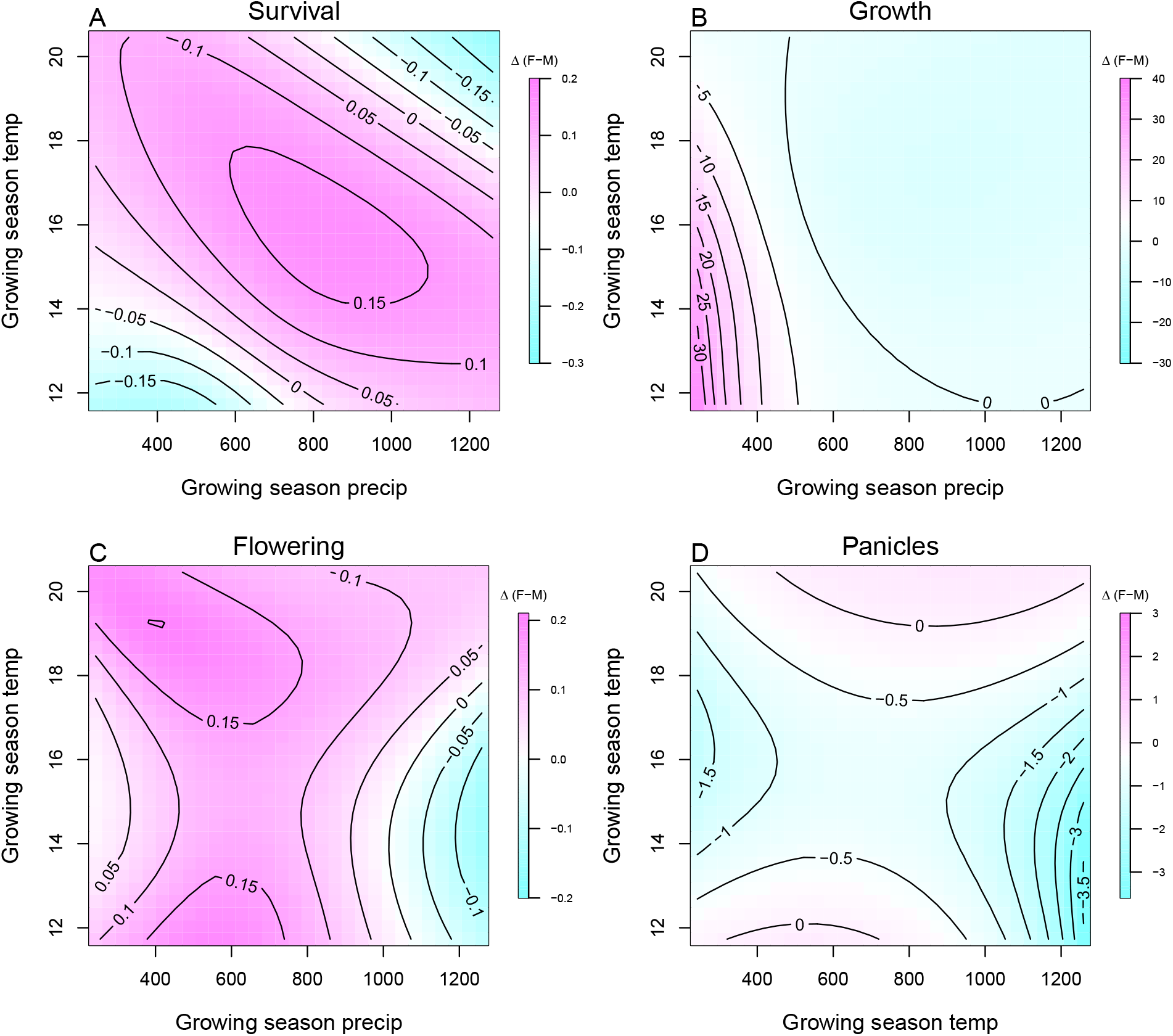
Predicted difference in demographic response to climate across species range between female and male. Contours show predicted value of that difference conditional on precipitation and temperature of growing season

**Figure S-7:**
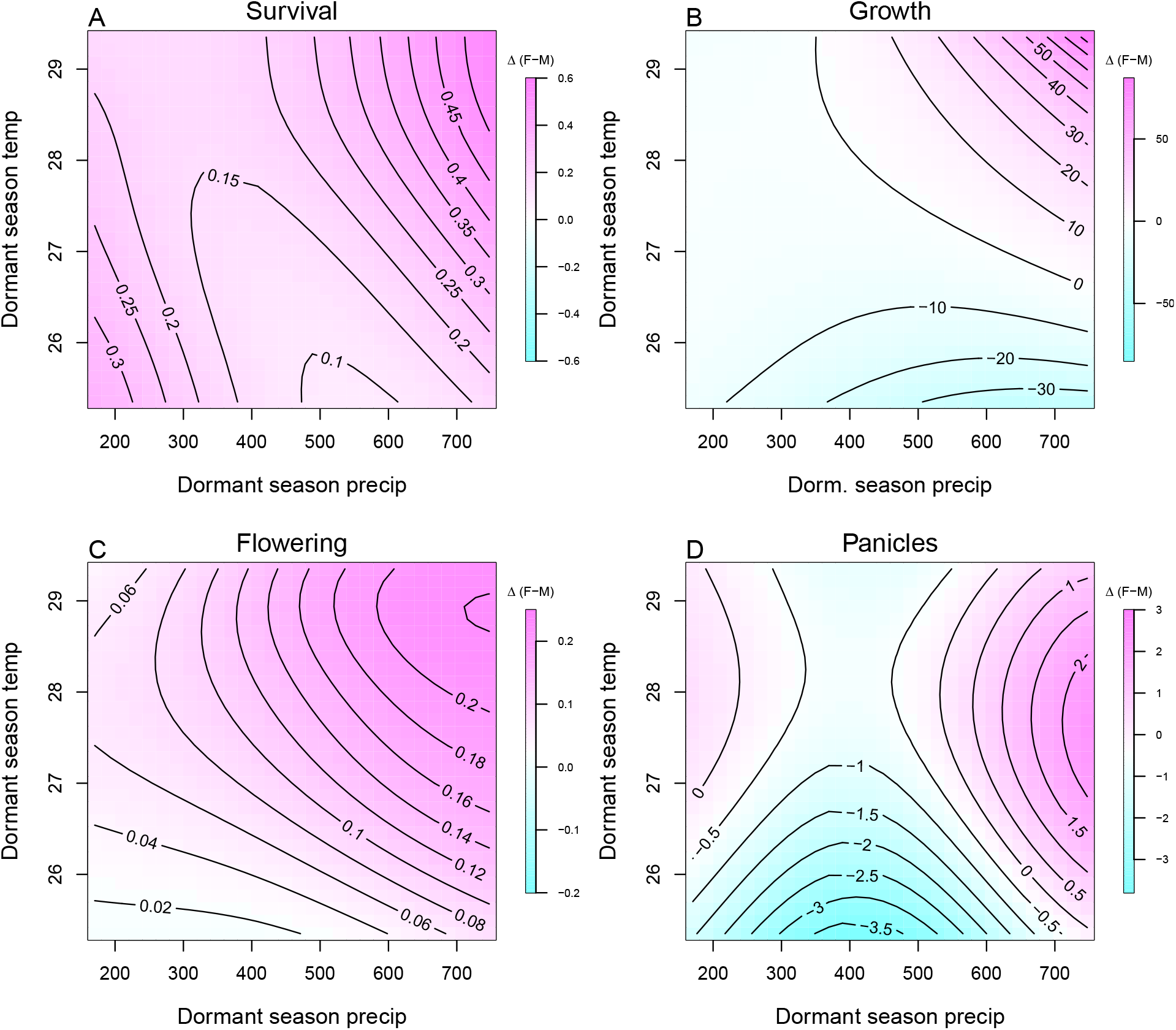
Predicted difference in demographic response to climate across species range between female and male. Contours show predicted value of that difference conditional on precipitation and temperature of the dormant season

**Figure S-8:**
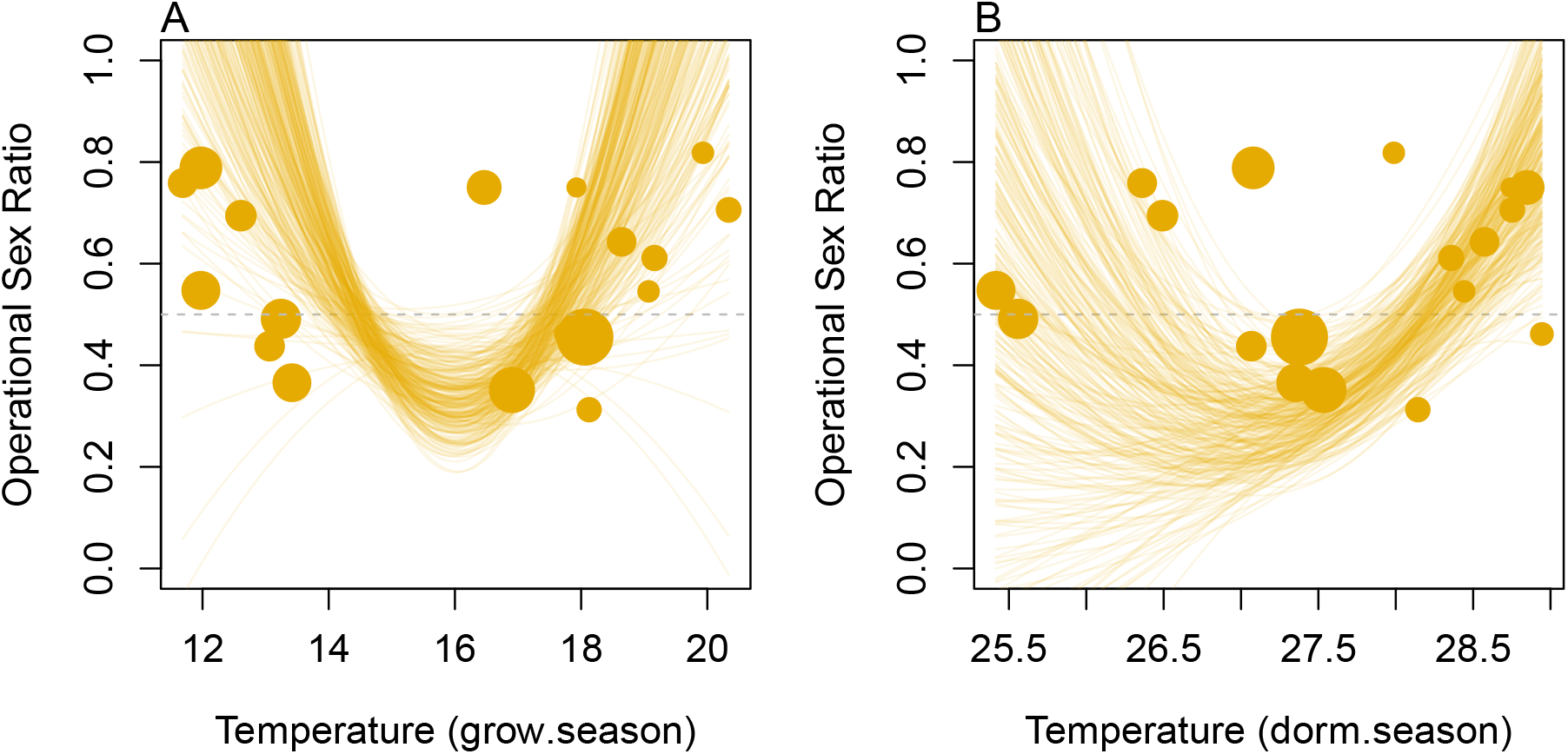
Significant Operational Sex Ratio response across climate gradient. (A, B) Proportion of panicles that were females across temperature of the growing and dormant season

**Figure S-9:**
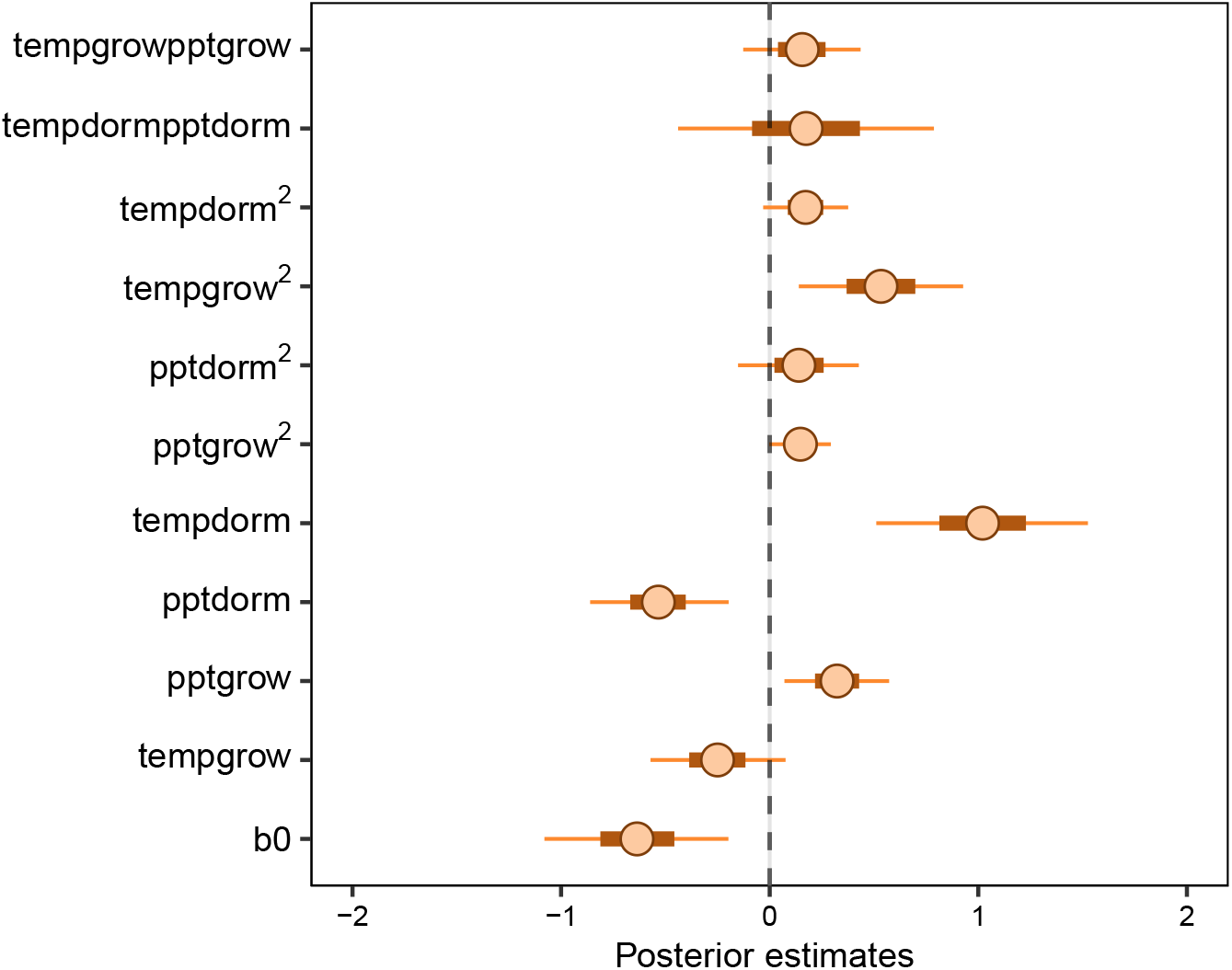
Mean parameter values and 95% credible intervals of the posterior probability distributions for climate drivers of operational sex ratio (female fraction of total panicles) across common garden years and sites. pptgrow is the precipitation of growing season, Tempgrow is the temperature of growing season, pptdorm is the temperature of dormant season, Tempdorm is the temperature of dormant season.

**Figure S-10:**
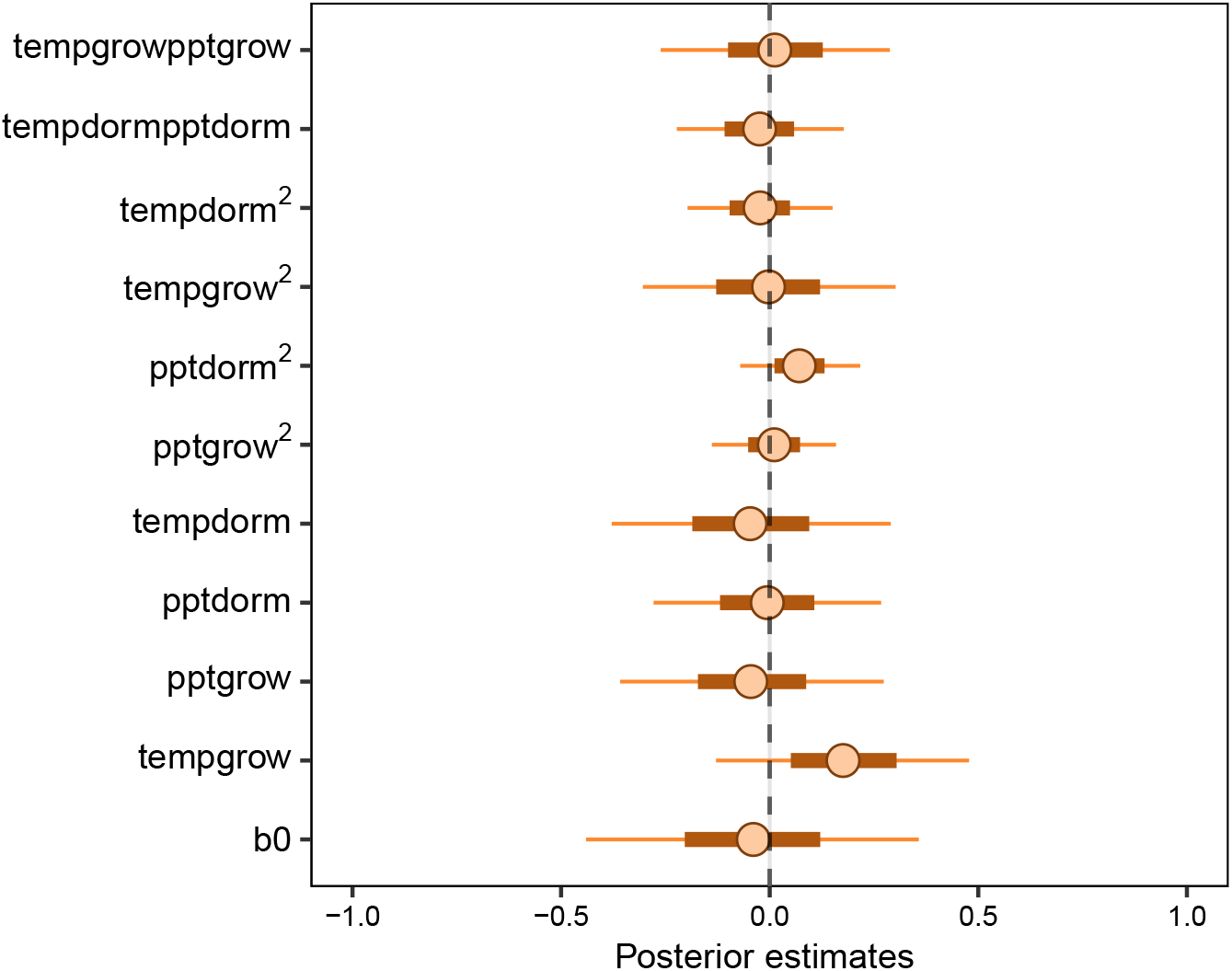
Mean parameter values and 95% credible intervals of the posterior probability distributions for climate drivers of sex ratio (female fraction of the populations) across common garden years and sites. pptgrow is the precipitation of growing season, Tempgrow is the temperature of growing season, pptdorm is the temperature of dormant season, Tempdorm is the temperature of dormant season.

**Figure S-11:**
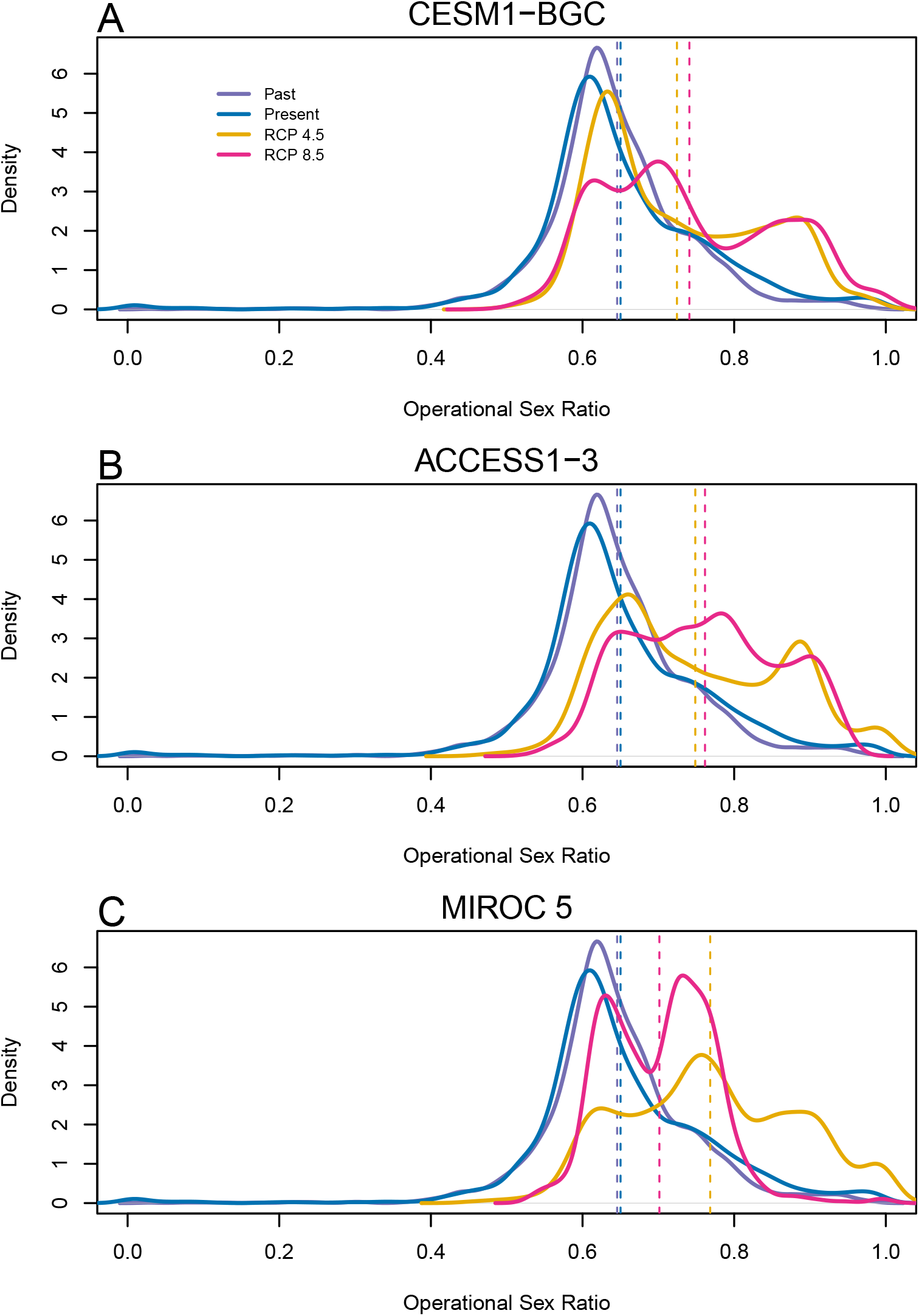
Change in Operational Sex Ratio (proportion of female) over time (past, present, future). Future projections were based on A) MIROC 5, B) CES, C) ACC. The means for each model are shown as vertical dashed lines.

**Figure S-12:**
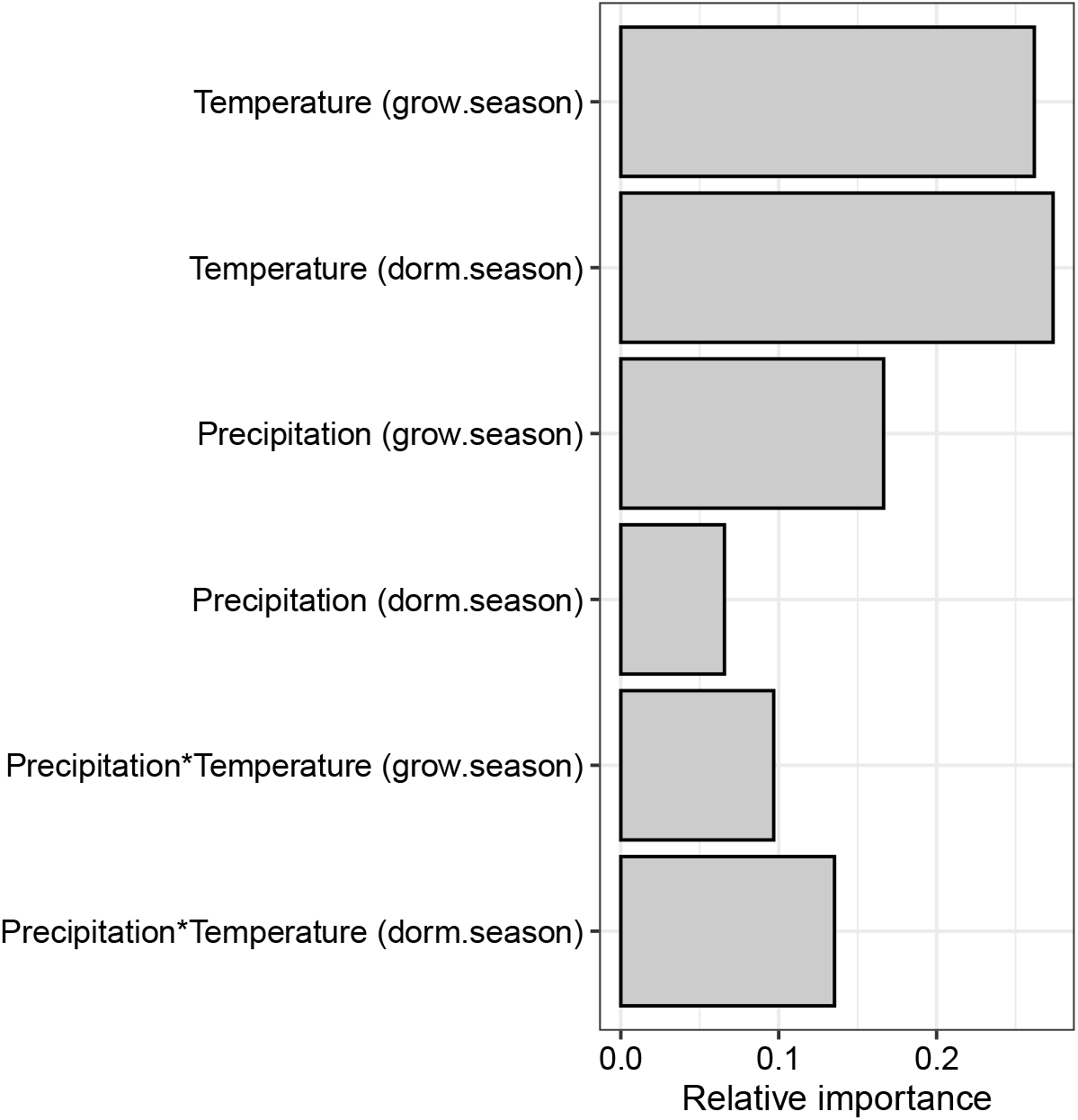
Life Table Response Experiment: The bar represent the relative importance of each predictors.

**Figure S-13:**
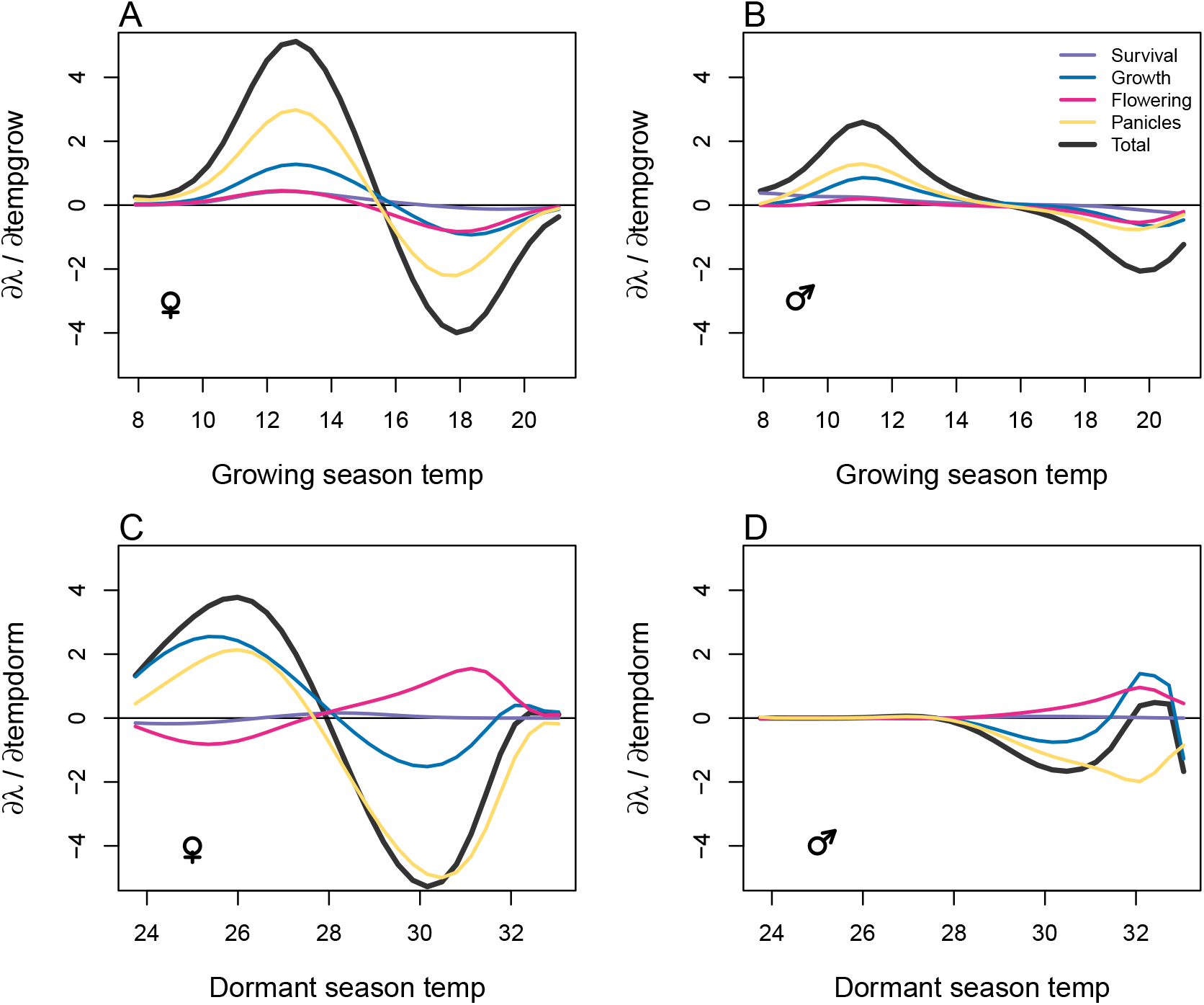
Life table response experiment decomposition of the sensitivity of *λ* to seasonal climate into additive vital rate contributions of males and females based on posterior mean parameter estimates. (A) Temperature of growing season (contribution of female), (B) Temperature of growing season (contribution of male), (C) Temperature of dormant season (contribution of female) and (D) Temperature of dormant season (contribution of male).

**Figure S-14:**
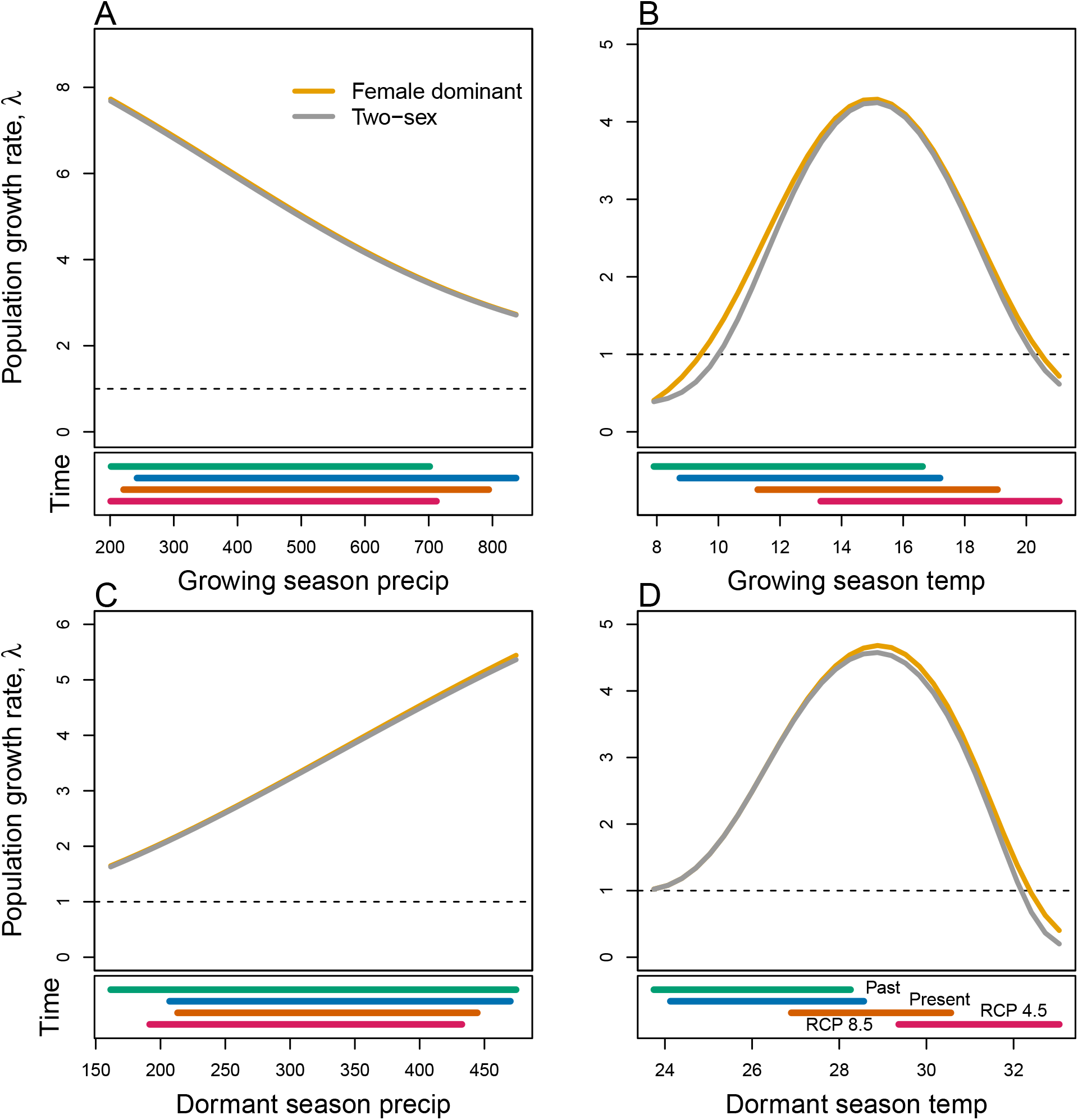
Predicted population growth rate (*λ*) in different ranges of climate. (A) Precipitation of the growing season, (B) Temperature of the growing season, (C) Precipitation of the dormant season, (D) Temperature of the dormant season. The grey curve shows prediction by the two-sex matrix projection model that incorporates sex-specific demographic responses to climate with sex ratio dependent seed fertilization. The orange curve represents the prediction by the female dominant matrix projection model. The dashed horizontal line indicates the limit of population viability (*λ* = 1). Lower panels below each data panel shows ranges of climate values for different time period (past climate, present and future climates). For future climate, we show a Representation Concentration Pathways (RCP) 4.5 and 8.5. Values population viability of (*λ*) are derived from the mean climate variables across 4 GCMs (MIROC5, ACCESS1-3, CESM1-BGC, CMCC-CM).

**Figure S-15:**
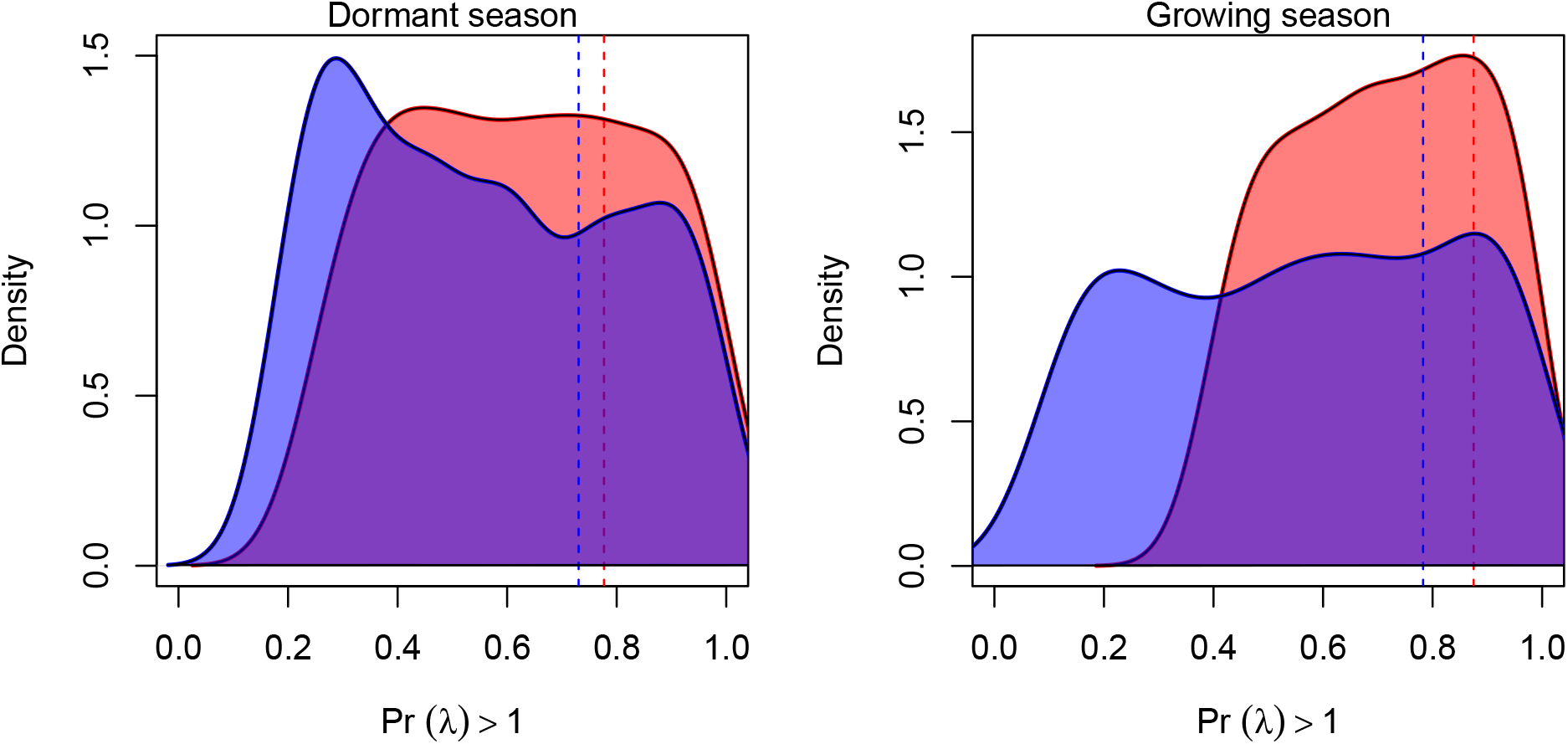
Assessment of the statistical difference between the two-sex models and the female dominant model for the dormant and growing season. Plots show the density of Pr (*λ*> 1) values for female dominant (pink) and two-sex models (violet) for each season. The means for each model are shown as vertical dashed lines.

**Figure S-16:**
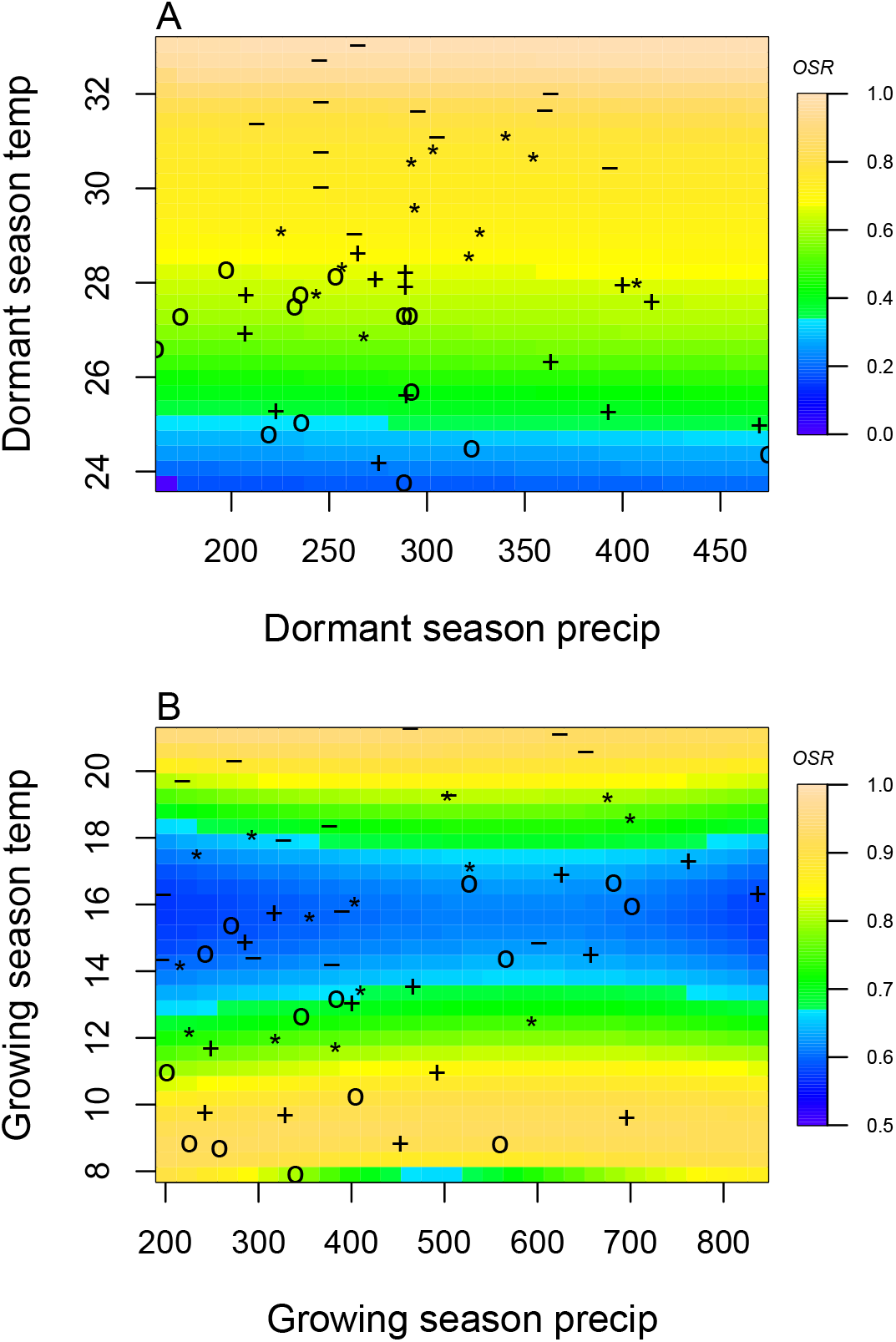
A two-dimensional representation of the Operational Sex Ratio (OSR) over time (past, present and future climate conditions). OSR represents the proportion of females. Contours show predicted values of OSR conditional on precipitation and temperature of the dormant and growing season. Operational Sex Ratio during the dormant season for the two sex model (A), Operational Sex Ratio during the growing season for the two sex model (B). “**o**”: Past, “**+**”: Current,”**\***”: RCP 4.5,”**-**”: RCP 8.5.

**Figure S-17:**
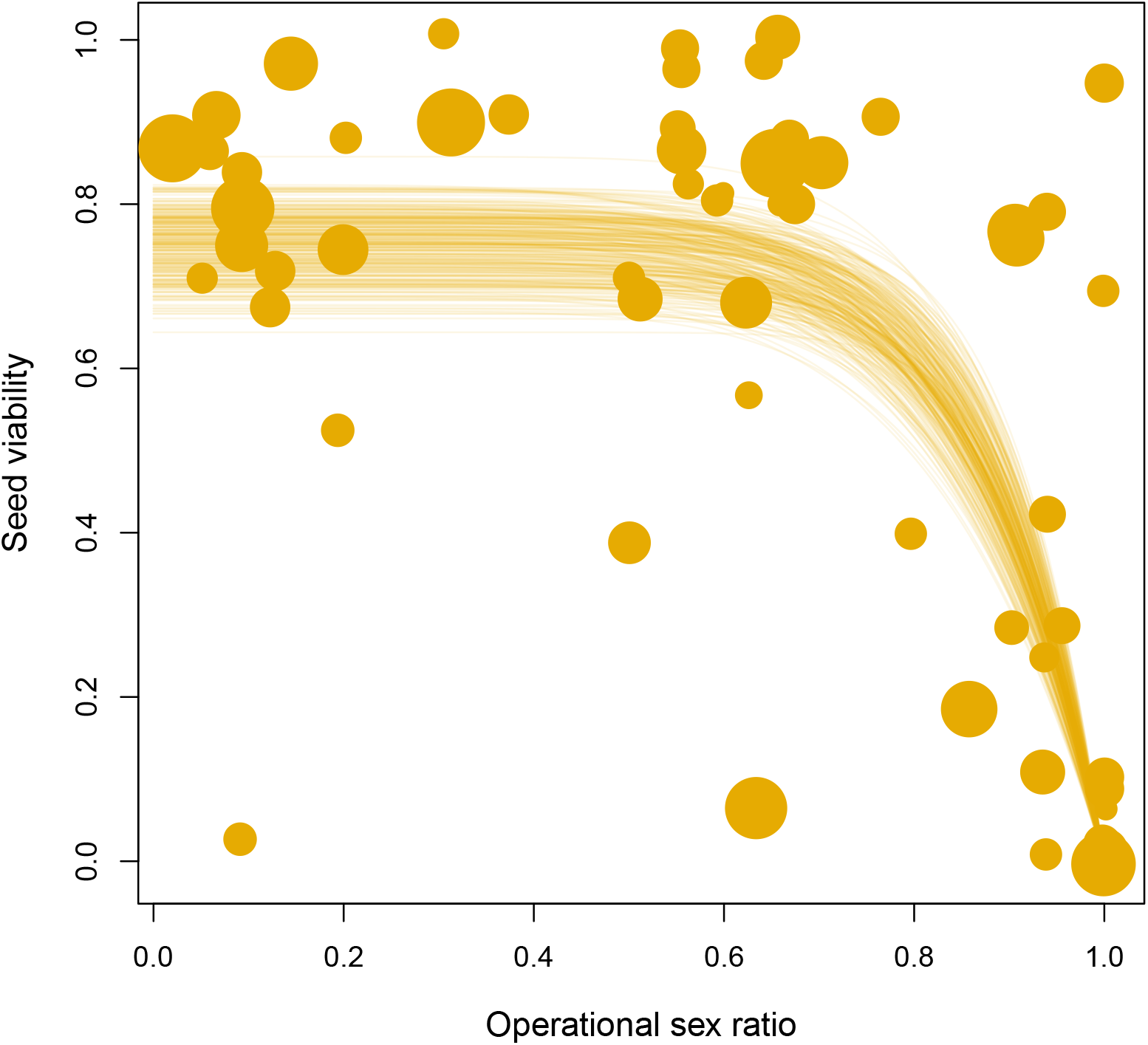
Seed fertilization success as a function of operational sex ratio (fraction of panicles that are female) in experimental populations. Circles show data from tetrazolium assays of seed viability; circle size is proportional to the number of seeds tested (minimum: 14; maximum: 57). Lines show model predictions for 300 samples from the posterior distribution of parameter estimate

**Figure S-18:**
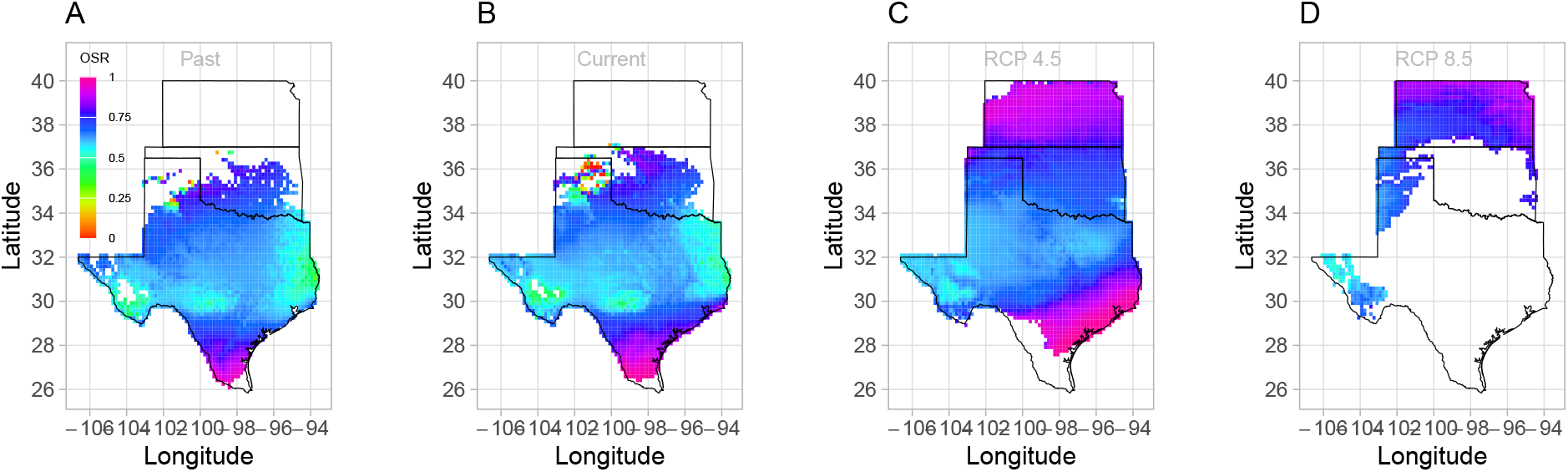
Projection of Operational Sex Ratio (proportion of female panicle) over time (past, present, future) across species range. Future projections were based on the CMCC-CM model.

**Figure S-19:**
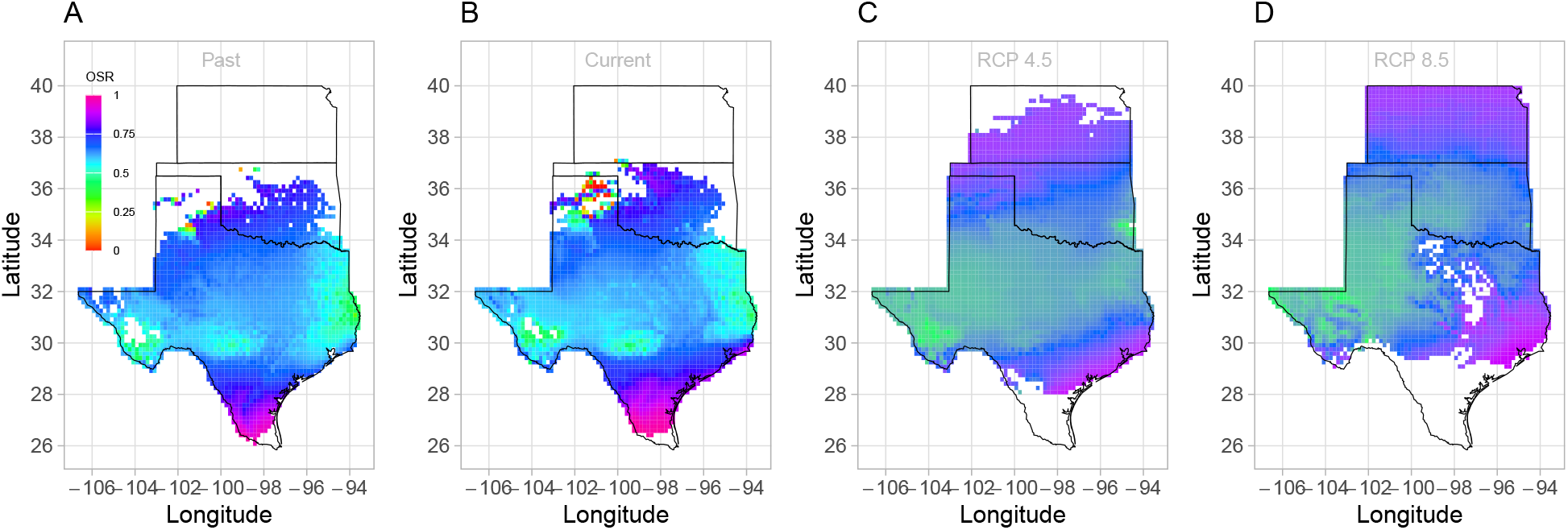
Projection of in Operational Sex Ratio (proportion of female panicle) over time (past, present, future) across species range. Future projections were based on the CES model.

**Figure S-20:**
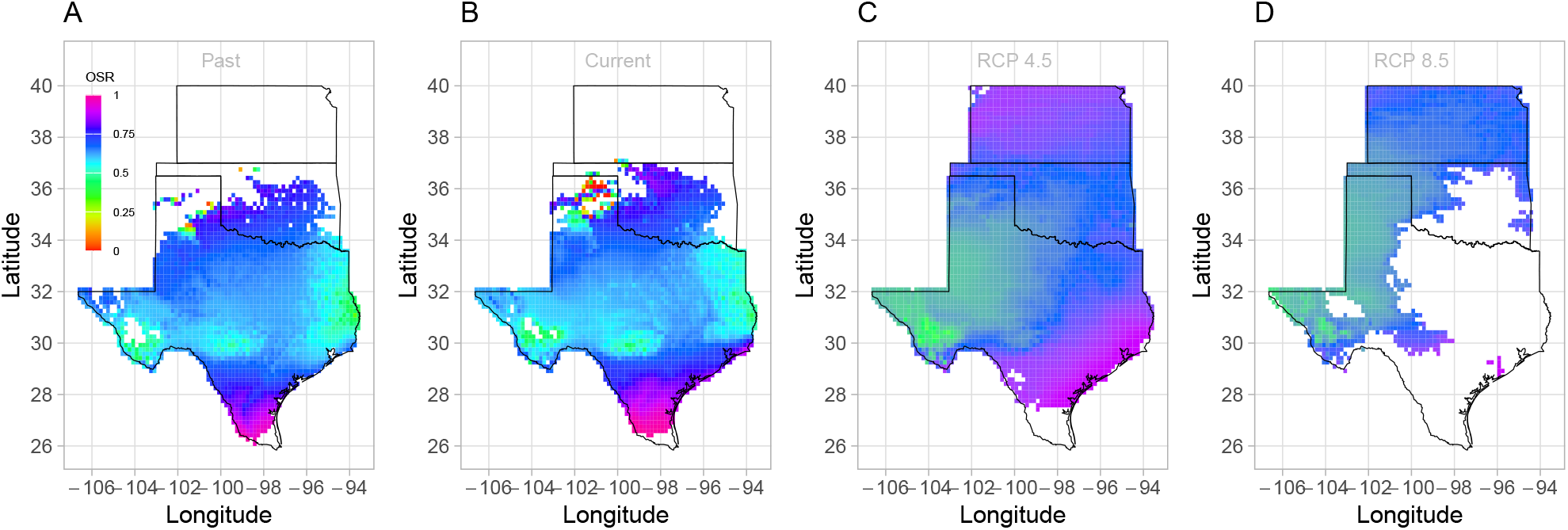
Projection of Operational Sex Ratio (proportion of female panicle) over time (past, present, future) across species range. Future projections were based on the MIROC model.

**Figure S-21:**
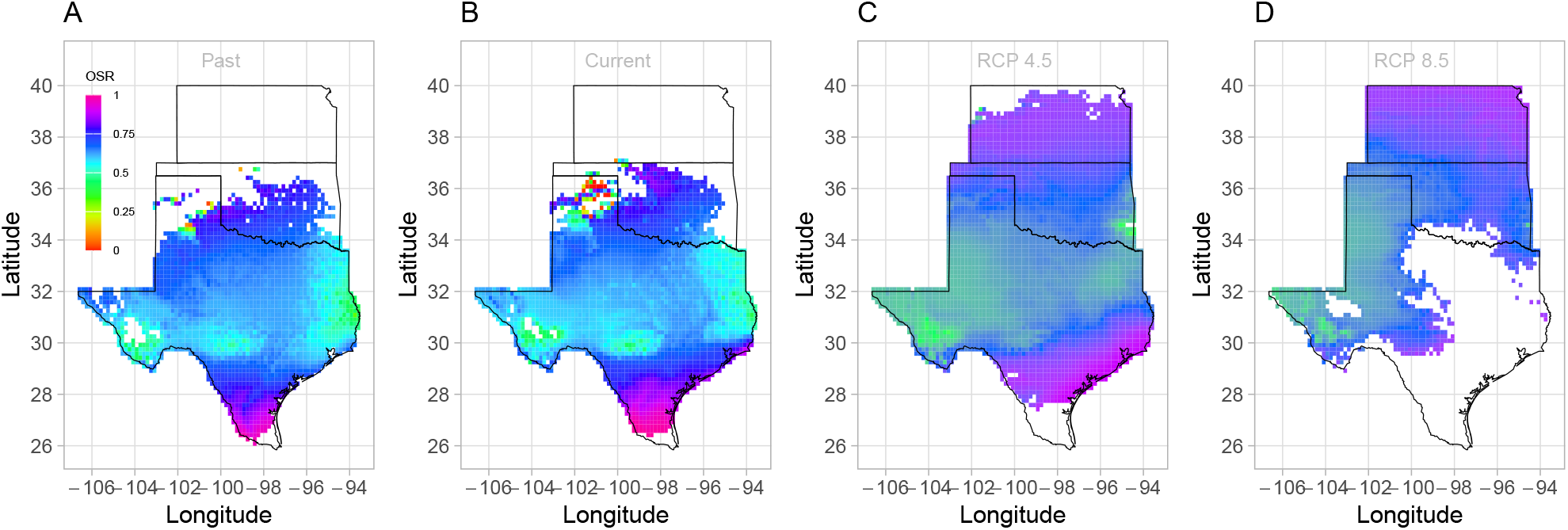
Projection of Operational Sex Ratio (proportion of female panicle) over time (past, present, future) across species range. Future projections were based on the ACCESS model. The mean sex ratio for each time period is shown as vertical dashed line.

**Figure S-22:**
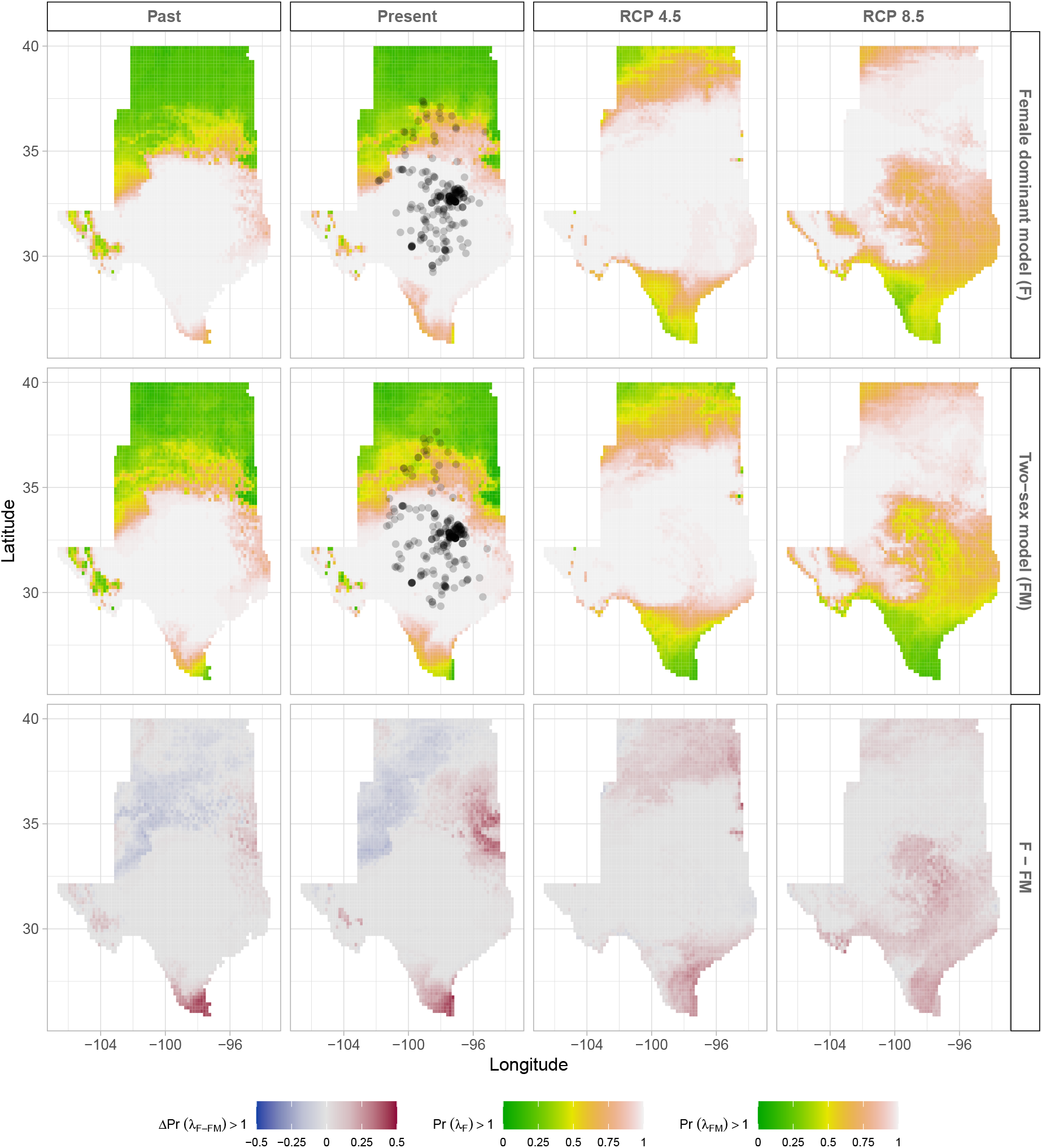
Climate change favors range shift toward the North edge of the current range. (A) Past, (B) Current, (C and D) Future predicted range shift based on the predicted probabilities of self-sustaining populations, Pr (*λ*> 1), using the two-sex model that incorporates sex-specific demographic responses to climate with sex ratio dependent seed fertilization. (E) Past, (F) Current, (G and F) Future predicted range shift based on the predicted probabilities of self-sustaining populations, Pr (*λ* > 1), using the female dominant model. Future projections were based on the CESM1-BGC model. The black dots on panel B and F indicate all known presence points collected from GBIF from 1990 to 2019, which corresponds to the current condition in our prediction. The occurrences of GBIFs are distributed in with higher population fitness habitat Pr (*λ*> 1), confirming that our study approach can reasonably predict range shifts.

**Figure S-23:**
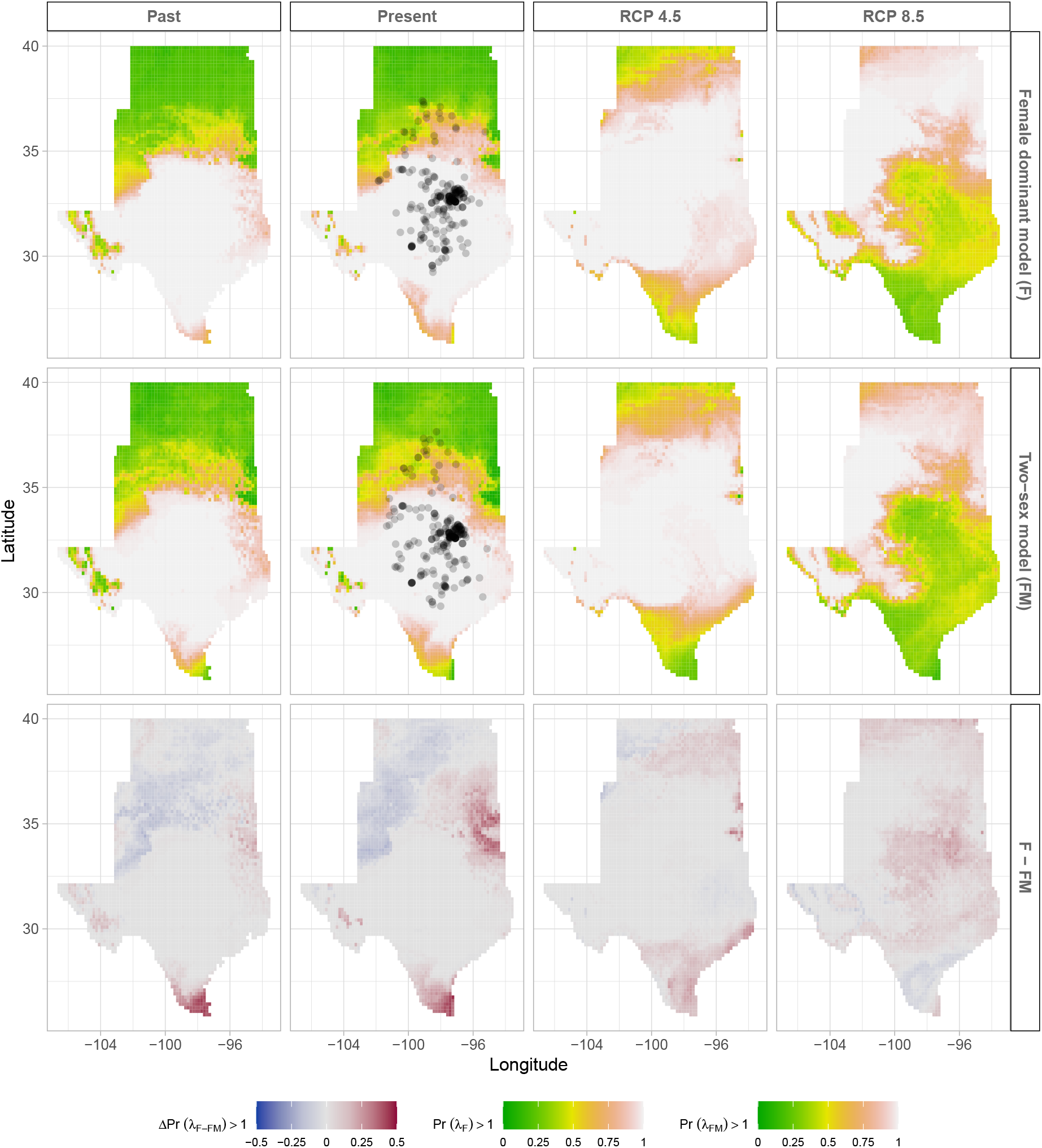
Climate change favors range shift toward the North edge of the current range. (A) Past, (B) Current, (C and D) Future predicted range shift based on the predicted probabilities of self-sustaining populations, Pr (*λ*> 1), using the two-sex model that incorporates sex-specific demographic responses to climate with sex ratio dependent seed fertilization. (E) Past, (F) Current, (G and F) Future predicted range shift based on the predicted probabilities of self-sustaining populations, Pr (*λ*> 1), using the female dominant model. Future projections were based on the ACCESS model. The black dots on panel B and F indicate all known presence points collected from GBIF from 1990 to 2019, which corresponds to the current condition in our prediction. The occurrences of GBIFs are distributed in with higher population fitness habitat Pr (*λ*> 1), confirming that our study approach can reasonably predict range shifts.

**Figure S-24:**
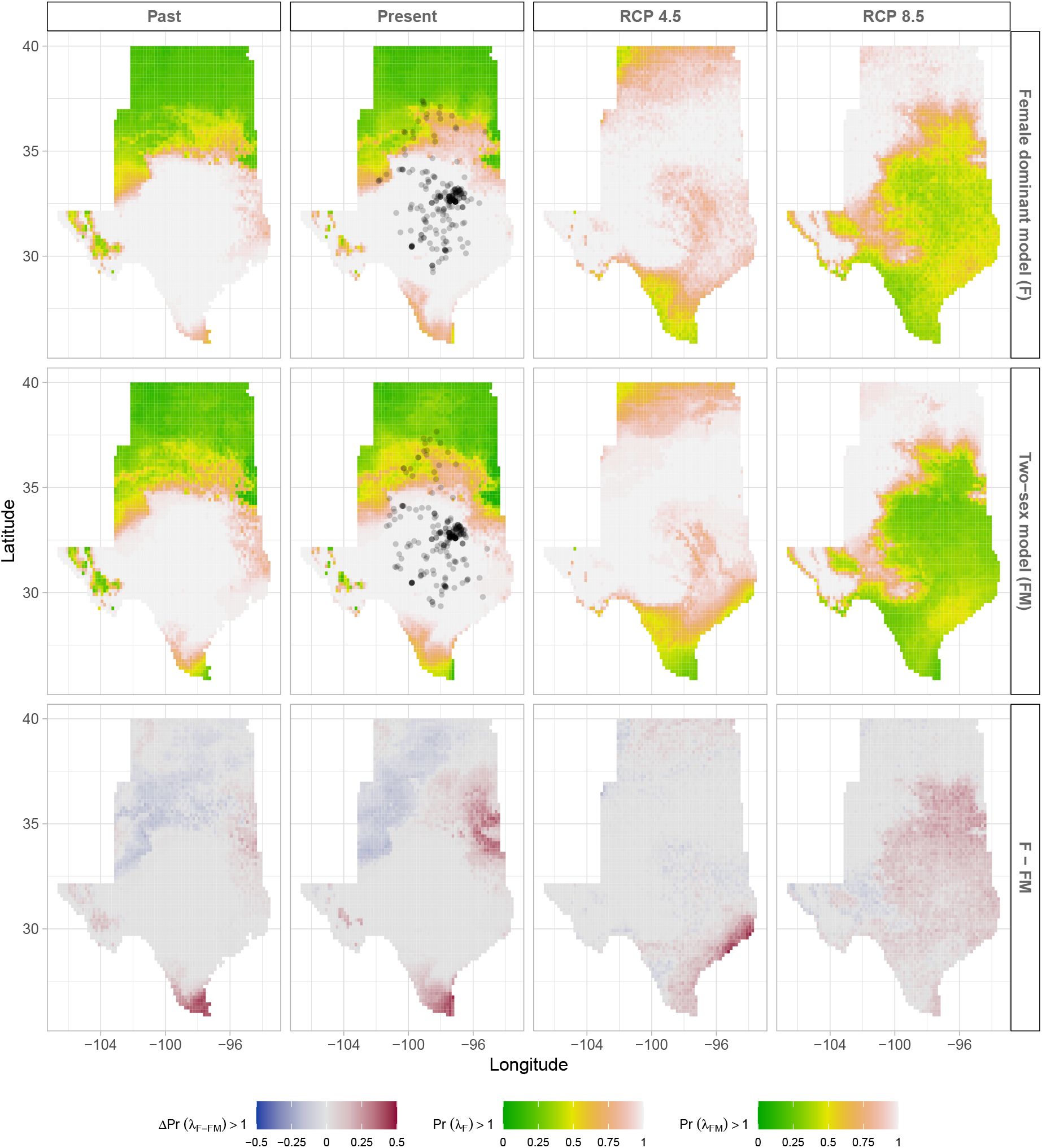
Climate change favors range shift toward the North edge of the current range. (A) Past, (B) Current, (C and D) Future predicted range shift based on the predicted probabilities of self-sustaining populations, Pr (*λ*> 1), using the two-sex model that incorporates sex-specific demographic responses to climate with sex ratio dependent seed fertilization. (E) Past, (F) Current, (G and F) Future predicted range shift based on the predicted probabilities of self-sustaining populations, Pr (*λ* > 1), using the female dominant model. Future projections were based on the MIROC5 model. The black dots on panel B and F indicate all known presence points collected from GBIF from 1990 to 2019, which corresponds to the current condition in our prediction. The occurrences of GBIFs are distributed in with higher population fitness habitat Pr (*λ*> 1), confirming that our study approach can reasonably predict range shifts.

**Figure S-25:**
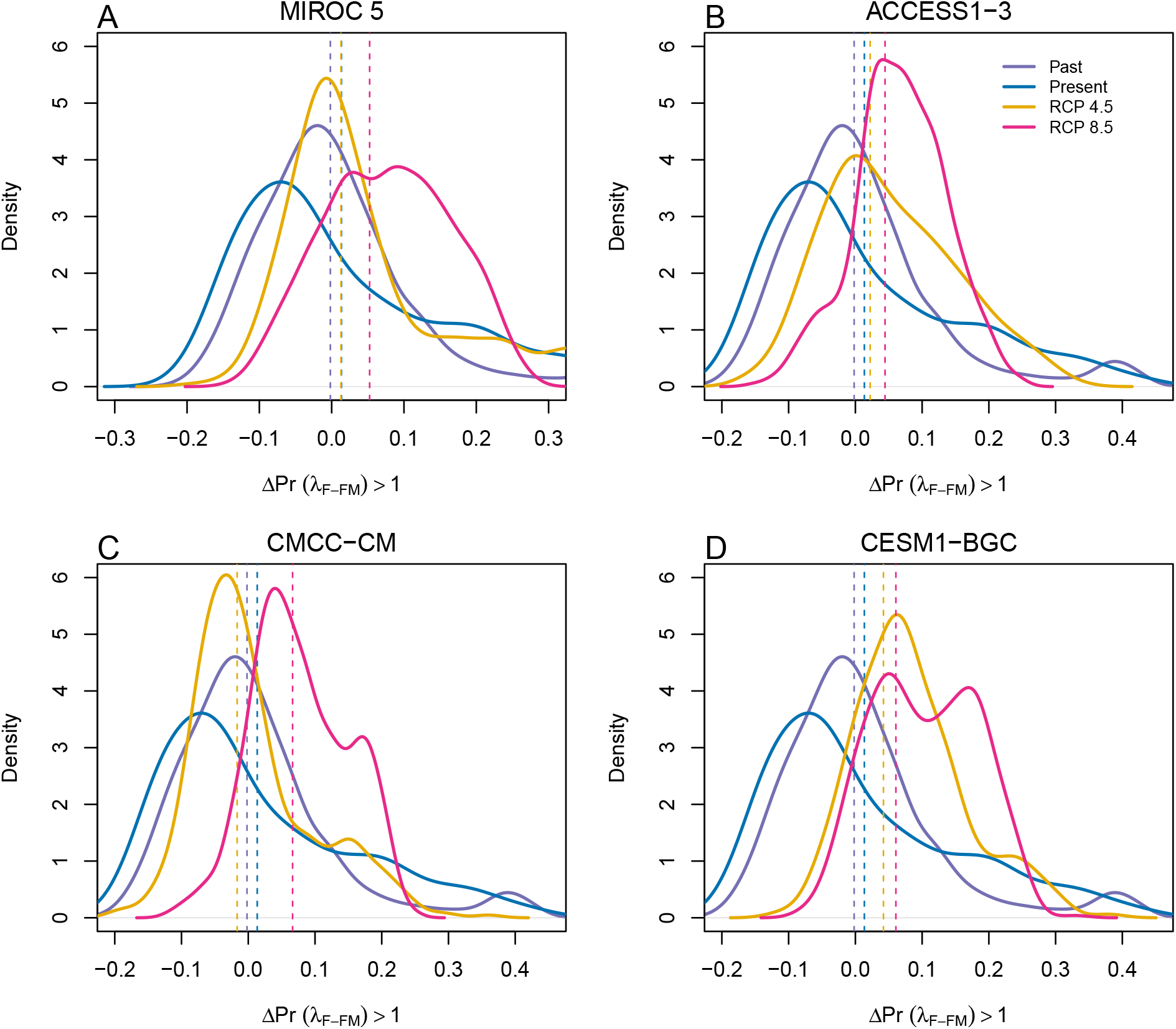
Assessment of the statistical difference between the two-sex models and the female dominant model across and beyond species range for 4 GCMs. The means for each model are shown as vertical dashed lines.

### S.2 Supporting Methods

#### S.2.1 Sex-specific demographic responses to climatic variation across common garden sites

Vital rate models were fit with the same linear predictors for the expected value (*µ*)(Eq.S.1): All vital rates were fit with second-degree polynomial functions to accommodate the possibility of hump-shaped relationships (reduced demographic performance at both extremes). We also included two-way interactions between sex and each climate driver and between temperature and precipitation within each season, and a three-way interaction between sex, temperature, and precipitation within each season. We modeled survival and flowering data with a Bernoulli distribution and the growth (tiller number) with a zero-truncated Poisson inverse Gaussian distribution. Fertility (panicle count conditional on flowering) was modeled as zero-truncated negative binomial. We used generic, weakly informative priors to fit coefficients for survival, growth, flowering models (*β*∼*N*(0, 1.5)) and random effect variances (*σ* ∼ *Gamma*(*γ*(0.1, 0.1)). We fit fertility model with also weakly informative priors for coefficients (*β*∼*N*(0, 0.15)). Different priors were used for fertility because the panicle model has a large number of parameters relative to the amount of available data (subset of our data) and because these specifics priors help prevent the model from overfitting. Each vital rate also includes normally distributed random effects for block-to-block variation (*ϕ*∼ *N*(0, *σ*_*block*_)), site to site variation (ν ∼ *N*(0, *σ*_*site*_)), and source-to-source variation that is related to the genetic provenence of the transplants used to establish the common garden (*ρ*∼*N*(0, *σ*_*source*_)).

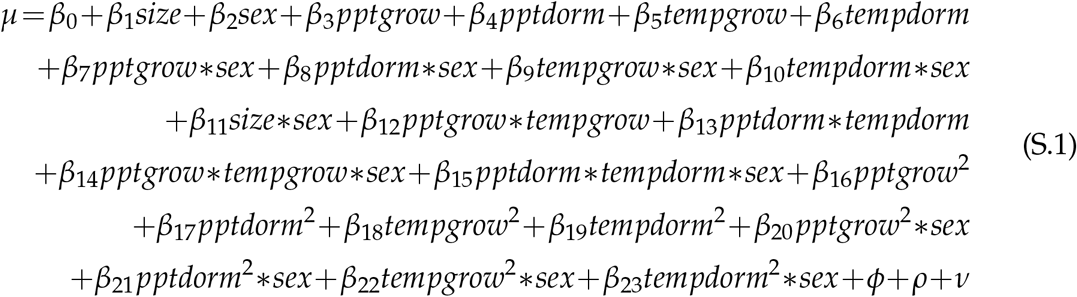

where *β*_0_ is the grand mean intercept, *β*_1_ is the size dependent slopes. *size* was on a natural logarithm scale. *β*_2_…*β*_13_ represent the climate dependent slopes. *β*_14_…*β*_23_ represent the sex-climate interaction slopes. *pptgrow* is the precipitation of the growing season, *tempgrow* is the temperature of the growing season, *pptdorm* is the precipitation of the dormant season, *tempdorm* is the temperature of the dormant season.

#### S.2.2 Sex ratio responses to climatic variation across common garden sites

To understand the impact of climatic variation across common garden sites on sex ratio, OSR and SR models using the same linear predictors for the expected value (ν)(Eq.S.2):

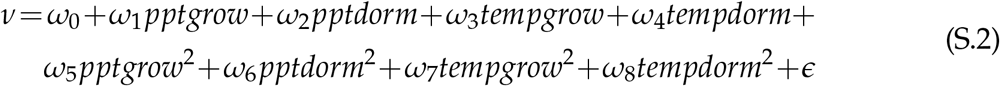

where *OSR* is the proportion of panicles that were female or proportion of female individuals in the experimental populations, c is the climate. *ω*_0_ is the intercept, *ω*_1_, *ω*_8_ are the climate dependent slopes. *ϵ* is error term.

We modeled the OSR and SR data with a Bernoulli distribution and used non informative priors for each coefficient (*ω*∼*N*(0, 100)).

#### S.2.3 Sex ratio experiment

To estimate the probability of seed viability, the germination rate and the effect of sex-ratio variation on female reproductive success, we conducted a sex-ratio experiment at one site near the center of the range to estimate the effect of sex-ratio variation on female reproductive success. The details of the experiment are provided in Compagnoni et al. (2017) and Miller and Compagnoni (2022b). Here we provide a summary of the experiment. We established 124 experimental populations in plots measuring 0.4 × 0.4m and separated by at least 15m from each other. We varied population density (1-48 plants/plot) and sex ratio (0%-100% female) across the experimental populations, and we replicated 34 combinations of density and sex ratio. We collected panicles from a subset of females in each plot and recorded the number of seeds in each panicle. We assessed reproductive success (seeds fertilized) using greenhouse-based germination and trazolium-based seed viability assays. Seed viability was modeled with a binomial distribution where the probability of viability (*v*) was given by:

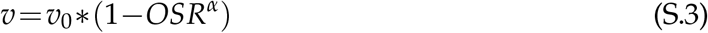

where *OSR* is the proportion of panicles that were female in the experimental populations. *α* is the parameter that control for how viability declines with increasing female bias. Further, germination rate was modeled using a binomial distribution to model the germination data from greenhouse trials. Given that germination was conditional on seed viability, the probability of success was given by the product *v*\**g*, where *v* is a function of *OSR* (Eq. S.3) and *g* is assumed to be constant.

## References

Angert, A. L., Bontrager, M. G., and Ågren, J. (2020). What do we really know about adaptation at range edges? Annual Review of Ecology, Evolution, and Systematics, 51(1):341–361.

Bawa, K. S. (1980). Evolution of dioecy in flowering plants. Annual review of ecology and systematics, 11:15–39.

Bertrand, R., Lenoir, J., Piedallu, C., Riofrío-Dillon, G., De Ruffray, P., Vidal, C., Pierrat, J.-C., and Gégout, J.-C. (2011). Changes in plant community composition lag behind climate warming in lowland forests. Nature, 479(7374):517–520.

Bruijning, M., Visser, M. D., Muller-Landau, H. C., Wright, S. J., Comita, L. S., Hubbell, S. P., de Kroon, H., and Jongejans, E. (2017). Surviving in a cosexual world: A cost-benefit analysis of dioecy in tropical trees. The American Naturalist, 189(3):297–314.

Bürli, S., Pannell, J. R., and Tonnabel, J. (2022). Environmental variation in sex ratios and sexual dimorphism in three wind-pollinated dioecious plant species. Oikos, 2022(6):e08651.

Caswell, H. (1989). Analysis of life table response experiments i. decomposition of effects on population growth rate. Ecological Modelling, 46(3-4):221–237.

Caswell, H. (2000). Matrix population models, volume 1. Sinauer Sunderland, MA.

Cipollini, M. L. and Whigham, D. F. (1994). Sexual dimorphism and cost of reproduction in the dioecious shrub lindera benzoin (lauraceae). American Journal of Botany, 81(1):65–75.

Cleland, E. E., Chuine, I., Menzel, A., Mooney, H. A., and Schwartz, M. D. (2007). Shifting plant phenology in response to global change. Trends in ecology & evolution, 22(7):357–365.

Compagnoni, A., Steigman, K., and Miller, T. E. (2017). Can’t live with them, can’t live without them? balancing mating and competition in two-sex populations. Proceedings of the Royal Society B: Biological Sciences, 284(1865):20171999.

Czachura, K. and Miller, T. E. (2020). Demographic back-casting reveals that subtle dimensions of climate change have strong effects on population viability. Journal of Ecology, 108(6):2557–2570.

Dahlgren, J. P., Bengtsson, K., and Ehrlén, J. (2016). The demography of climate-driven and density-regulated population dynamics in a perennial plant. Ecology, 97(4):899–907.

Davis, M. B. and Shaw, R. G. (2001). Range shifts and adaptive responses to quaternary climate change. Science, 292(5517):673–679.

Dietze, M. C., Fox, A., Beck-Johnson, L. M., Betancourt, J. L., Hooten, M. B., Jarnevich, C. S., Keitt, T. H., Kenney, M. A., Laney, C. M., Larsen, L. G., et al. (2018). Iterative near-term ecological forecasting: Needs, opportunities, and challenges. Proceedings of the National Academy of Sciences, 115(7):1424–1432.

Diez, J. M., Giladi, I., Warren, R., and Pulliam, H. R. (2014). Probabilistic and spatially variable niches inferred from demography. Journal of ecology, 102(2):544–554.

Eberhart-Phillips, L. J., Küpper, C., Miller, T. E., Cruz-López, M., Maher, K. H., Dos Remedios, N., Stoffel, M. A., Hoffman, J. I., Krüger, O., and Székely, T. (2017). Sex-specific early survival drives adult sex ratio bias in snowy plovers and impacts mating system and population growth. Proceedings of the National Academy of Sciences, 114(27):E5474–E5481.

Ehrlén, J. and Morris, W. F. (2015). Predicting changes in the distribution and abundance of species under environmental change. Ecology letters, 18(3):303–314.

Elderd, B. D. and Miller, T. E. (2016). Quantifying demographic uncertainty: Bayesian methods for integral projection models. Ecological Monographs, 86(1):125–144.

Ellis, R. P., Davison, W., Queirós, A. M., Kroeker, K. J., Calosi, P., Dupont, S., Spicer, J. I., Wilson, R. W., Widdicombe, S., and Urbina, M. A. (2017). Does sex really matter? explaining intraspecies variation in ocean acidification responses. Biology letters, 13(2):20160761.

Ellner, S. P., Adler, P. B., Childs, D. Z., Hooker, G., Miller, T. E., and Rees, M. (2022). A critical comparison of integral projection and matrix projection models for demographic analysis: Comment. Ecology.

Ellner, S. P., Childs, D. Z., Rees, M., et al. (2016). Data-driven modelling of structured populations. A practical guide to the Integral Projection Model. Cham: Sprimger.

Evans, M. E., Merow, C., Record, S., McMahon, S. M., and Enquist, B. J. (2016). Towards process-based range modeling of many species. Trends in Ecology & Evolution, 31(11):860–871.

Field, D. L., Pickup, M., and Barrett, S. C. (2013). Comparative analyses of sex-ratio variation in dioecious flowering plants. Evolution, 67(3):661–672.

Gamelon, M., Grøtan, V., Nilsson, A. L., Engen, S., Hurrell, J. W., Jerstad, K., Phillips, A. S., Røstad, O. W., Slagsvold, T., Walseng, B., et al. (2017). Interactions between demography and environmental effects are important determinants of population dynamics. Science Advances, 3(2):e1602298.

Gerber, L. R. and White, E. R. (2014). Two-sex matrix models in assessing population viability: when do male dynamics matter? Journal of Applied Ecology, 51(1):270–278.

Gilbert, K. J., Sharp, N. P., Angert, A. L., Conte, G. L., Draghi, J. A., Guillaume, F., Hargreaves, A. L., Matthey-Doret, R., and Whitlock, M. C. (2017). Local adaptation interacts with expansion load during range expansion: maladaptation reduces expansion load. The American Naturalist, 189(4):368–380.

Gissi, E., Bowyer, R. T., and Bleich, V. C. (2024). Sex-based differences affect conservation. Science, 384(6702):1309–1310.

Gissi, E., Schiebinger, L., Hadly, E. A., Crowder, L. B., Santoleri, R., and Micheli, F. (2023). Exploring climate-induced sex-based differences in aquatic and terrestrial ecosystems to mitigate biodiversity loss. nature communications, 14(1):4787.

Hernández, C. M., Ellner, S. P., Adler, P. B., Hooker, G., and Snyder, R. E. (2023). An exact version of life table response experiment analysis, and the r package exactltre. Methods in Ecology and Evolution, 14(3):939–951.

Hitchcock, A. S. (1971). Manual of the grasses of the United States, volume 2. Courier Corporation.

Hultine, K. R., Grady, K. C., Wood, T. E., Shuster, S. M., Stella, J. C., and Whitham, T. G. (2016). Climate change perils for dioecious plant species. Nature Plants, 2(8):1–8.

Iler, A. M., Compagnoni, A., Inouye, D. W., Williams, J. L., CaraDonna, P. J., Anderson, A., and Miller, T. E. (2019). Reproductive losses due to climate change-induced earlier flowering are not the primary threat to plant population viability in a perennial herb. Journal of Ecology, 107(4):1931–1943.

Intergovernmental Panel On Climate Change (Ipcc) (2023). Climate Change 2022 – Impacts, Adaptation and Vulnerability: Working Group II Contribution to the Sixth Assessment Report of the Intergovernmental Panel on Climate Change. Cambridge University Press, 1 edition.

Jenouvrier, S., Holland, M., Stroeve, J., Barbraud, C., Weimerskirch, H., Serreze, M., and Caswell, H. (2012). Effects of climate change on an emperor penguin population: analysis of coupled demographic and climate models. Global Change Biology, 18(9):2756–2770.

Jones, M. H., Macdonald, S. E., and Henry, G. H. (1999). Sex-and habitat-specific responses of a high arctic willow, salix arctica, to experimental climate change. Oikos, pages 129–138.

Karger, D. N., Conrad, O., Böhner, J., Kawohl, T., Kreft, H., Soria-Auza, R. W., Zimmermann, N. E., Linder, H. P., and Kessler, M. (2017). Climatologies at high resolution for the earth’s land surface areas. Scientific data, 4(1):1–20.

Kindiger, B. (2004). Interspecific hybrids of poa arachnifera× poa secunda. Journal of New Seeds, 6(1):1–26.

Konapala, G., Mishra, A. K., Wada, Y., and Mann, M. E. (2020). Climate change will affect global water availability through compounding changes in seasonal precipitation and evaporation. Nature communications, 11(1):3044.

Lee-Yaw, J. A., Kharouba, H. M., Bontrager, M., Mahony, C., Csergő, A. M., Noreen, A. M., Li, Q., Schuster, R., and Angert, A. L. (2016). A synthesis of transplant experiments and ecological niche models suggests that range limits are often niche limits. Ecology letters, 19(6):710–722.

Liaw, A., Wiener, M., et al. (2002). Classification and regression by randomforest. R news, 2(3):18–22.

Louthan, A. M., Keighron, M., Kiekebusch, E., Cayton, H., Terando, A., and Morris, W. F. (2022). Climate change weakens the impact of disturbance interval on the growth rate of natural populations of venus flytrap. Ecological Monographs, 92(4):e1528.

Lynch, H. J., Rhainds, M., Calabrese, J. M., Cantrell, S., Cosner, C., and Fagan, W. F. (2014). How climate extremes—not means—define a species’ geographic range boundary via a demographic tipping point. Ecological Monographs, 84(1):131–149.

Margaret EK, E., Sharmila MN, D., Heilman, K. A., Tipton, J. R., DeRose, R. J., Stefan, K., Schultz, E. L., and Shaw, J. D. (2023). The trailing edge is everywhere: tree rings reveal the transient risk of extinction hidden inside climate envelope forecasts. Technical report, Los Alamos National Laboratory (LANL), Los Alamos, NM (United States).

McLean, N., Lawson, C. R., Leech, D. I., and van de Pol, M. (2016). Predicting when climate-driven phenotypic change affects population dynamics. Ecology Letters, 19(6):595–608.

Merow, C., Bois, S. T., Allen, J. M., Xie, Y., and Silander Jr, J. A. (2017). Climate change both facilitates and inhibits invasive plant ranges in new england. Proceedings of the National Academy of Sciences, 114(16):E3276–E3284.

Miller, T. and Compagnoni, A. (2022a). Data from: Two-sex demography, sexual niche differentiation, and the geographic range limits of texas bluegrass (Poa arachnifera). American Naturalist, Dryad Digital Repository,. 10.5061/dryad.kkwh70s5x.

Miller, T. E. and Compagnoni, A. (2022b). Two-sex demography, sexual niche differentiation, and the geographic range limits of texas bluegrass (poa arachnifera). The American Naturalist, 200(1):17–31.

Miller, T. E., Shaw, A. K., Inouye, B. D., and Neubert, M. G. (2011). Sex-biased dispersal and the speed of two-sex invasions. The American Naturalist, 177(5):549–561.

Milner-Gulland, E. (1994). A population model for the management of the saiga antelope. Journal of Applied Ecology, pages 25–39.

Molnár, P. K., Derocher, A. E., Thiemann, G. W., and Lewis, M. A. (2010). Predicting survival, reproduction and abundance of polar bears under climate change. Biological Conservation, 143(7):1612–1622.

Morrison, C. A., Robinson, R. A., Clark, J. A., and Gill, J. A. (2016). Causes and consequences of spatial variation in sex ratios in a declining bird species. Journal of Animal Ecology, 85(5):1298–1306.

Pagel, J., Treurnicht, M., Bond, W. J., Kraaij, T., Nottebrock, H., Schutte-Vlok, A., Tonnabel, J., Esler, K. J., and Schurr, F. M. (2020). Mismatches between demographic niches and geographic distributions are strongest in poorly dispersed and highly persistent plant species. Proceedings of the National Academy of Sciences, 117(7):3663–3669.

Pease, C. M., Lande, R., and Bull, J. (1989). A model of population growth, dispersal and evolution in a changing environment. Ecology, 70(6):1657–1664.

Petry, W. K., Soule, J. D., Iler, A. M., Chicas-Mosier, A., Inouye, D. W., Miller, T. E., and Mooney, K. A. (2016). Sex-specific responses to climate change in plants alter population sex ratio and performance. Science, 353(6294):69–71.

Piironen, J. and Vehtari, A. (2017). Comparison of bayesian predictive methods for model selection. Statistics and Computing, 27:711–735.

Pottier, P., Burke, S., Drobniak, S. M., Lagisz, M., and Nakagawa, S. (2021). Sexual (in) equality? a meta-analysis of sex differences in thermal acclimation capacity across ectotherms. Functional Ecology, 35(12):2663–2678.

R Core Team (2023). R: A Language and Environment for Statistical Computing. R Foundation for Statistical Computing, Vienna, Austria.

Reed, P. B., Peterson, M. L., Pfeifer-Meister, L. E., Morris, W. F., Doak, D. F., Roy, B. A., Johnson, B. R., Bailes, G. T., Nelson, A. A., and Bridgham, S. D. (2021). Climate manipulations differentially affect plant population dynamics within versus beyond northern range limits. Journal of Ecology, 109(2):664–675.

Renganayaki, K., Jessup, R., Burson, B., Hussey, M., and Read, J. (2005). Identification of male-specific aflp markers in dioecious texas bluegrass. Crop science, 45(6):2529–2539.

Sanderson, B. M., Knutti, R., and Caldwell, P. (2015). A representative democracy to reduce interdependency in a multimodel ensemble. Journal of Climate, 28(13):5171–5194.

Sasaki, M., Hedberg, S., Richardson, K., and Dam, H. G. (2019). Complex interactions between local adaptation, phenotypic plasticity and sex affect vulnerability to warming in a widespread marine copepod. Royal Society open science, 6(3):182115.

Schultz, E. L., Hülsmann, L., Pillet, M. D., Hartig, F., Breshears, D. D., Record, S., Shaw, J. D., DeRose, R. J., Zuidema, P. A., and Evans, M. E. (2022). Climate-driven, but dynamic and complex? a reconciliation of competing hypotheses for species’ distributions. Ecology letters, 25(1):38–51.

Schwalm, C. R., Glendon, S., and Duffy, P. B. (2020). Rcp8. 5 tracks cumulative co2 emissions. Proceedings of the National Academy of Sciences, 117(33):19656–19657.

Schwinning, S., Lortie, C. J., Esque, T. C., and DeFalco, L. A. (2022). What common-garden experiments tell us about climate responses in plants.

Sexton, J. P., McIntyre, P. J., Angert, A. L., and Rice, K. J. (2009). Evolution and ecology of species range limits. Annu. Rev. Ecol. Evol. Syst., 40:415–436.

Shelton, A. O. (2010). The ecological and evolutionary drivers of female-biased sex ratios: two-sex models of perennial seagrasses. The American Naturalist, 175(3):302–315.

Sherry, R. A., Zhou, X., Gu, S., Arnone III, J. A., Schimel, D. S., Verburg, P. S., Wallace, L. L., and Luo, Y. (2007). Divergence of reproductive phenology under climate warming. Proceedings of the National Academy of Sciences, 104(1):198–202.

Smith, M. D., Wilkins, K. D., Holdrege, M. C., Wilfahrt, P., Collins, S. L., Knapp, A. K., Sala, O. E., Dukes, J. S., Phillips, R. P., Yahdjian, L., et al. (2024). Extreme drought impacts have been underestimated in grasslands and shrublands globally. Proceedings of the National Academy of Sciences, 121(4):e2309881120.

Stan Development Team (2023). RStan: the R interface to Stan. R package version 2.21.8.

Thomson, A. M., Calvin, K. V., Smith, S. J., Kyle, G. P., Volke, A., Patel, P., Delgado-Arias, S., Bond-Lamberty, B., Wise, M. A., Clarke, L. E., et al. (2011). Rcp4. 5: a pathway for stabilization of radiative forcing by 2100. Climatic change, 109:77–94.

Tognetti, R. (2012). Adaptation to climate change of dioecious plants: does gender balance matter? Tree Physiology, 32(11):1321–1324.

Williams, J. L., Jacquemyn, H., Ochocki, B. M., Brys, R., and Miller, T. E. (2015). Life history evolution under climate change and its influence on the population dynamics of a long-lived plant. Journal of Ecology, 103(4):798–808.

Zhao, H., Li, Y., Zhang, X., Korpelainen, H., and Li, C. (2012). Sex-related and stage-dependent source-to-sink transition in populus cathayana grown at elevated co 2 and elevated temperature. Tree Physiology, 32(11):1325–1338.

